# Nanofabricated ultraflexible electrodes for chronic, high-density neural recording and stimulation

**DOI:** 10.64898/2026.11.23.768676

**Authors:** Weinan Wang, Yuhang Ma, Pavlo Zolotavin, Liuyang Sun, Hanqing Wang, Kyu Hyun Lee, Kalman A. Katlowitz, Carter A. Kelly, Hanlin Zhu, Rongkang Yin, Yingchu Sun, Robin Kim, Luis De La Palma Piedras, Jiaao Zhang, Yuxuan Liu, Xiaorong Zhang, Elizabeth A. Mickiewicz, Philip Adenekan, Nicole R. Provenza, Ganesh Rao, Alla Y. Karpova, Benjamin Y. Hayden, Sameer A. Sheth, Loren M. Frank, Lan Luan, Chong Xie

## Abstract

Flexible neural probes enable stable, long-term recording and stimulation with minimal tissue disruption, but their electrode density and channel count remain limited. Here, we report an electron-beam lithography (EBL) manufacturing strategy for ultraflexible polymer probes that increases channel density by over tenfold, accommodating up to 512 channels within a cross-sectional area of 240 µm^2^. These high-density probes improve single-unit isolation, resolve functional connectivity, and enable months-long tracking of neuronal ensembles. Combined intracortical microstimulation and simultaneous recordings further enable spatially selective modulation and mapping of local microcircuits. We demonstrate these capabilities across diverse chronic animal models and establish translational feasibility through stable intraoperative recordings of lexical-semantic representations in human temporal cortex. Together, nanofabricated ultraflexible probes provide a versatile platform for longitudinal circuit investigations and high-density bidirectional brain–computer interfaces integrating neural recording with spatially selective stimulation.

## Introduction

Penetrating neural microelectrodes enable single-neuron recording with millisecond temporal resolutions by interfacing in close proximity to individual neurons. They have therefore become indispensable tools in systems neuroscience and promising platforms for investigating neurological disorders involving circuit dysfunctions. The need to resolve the dynamics of densely packed and interconnected neuronal populations has driven the decades-long pursuit of large-scale neural recording technologies^1–3^. Current state-of-the-art approaches predominantly rely on silicon microfabrication technologies, exemplified by Neuropixel probes^1,4,5^, and can simultaneously record hundreds to thousands of neurons at high spatial densities. However, despite these remarkable advances, a key limitation of silicon-based electrodes is the mechanical mismatch at the electrode-tissue interface. Following implantation, mechanical stresses between the rigid electrode and soft brain tissue induce short-term (second to hour) displacements that result in recording drifts^5,6^. Over longer timescales, persistent mechanical irritation often triggers tissue reactions, leading to degeneration of nearby neurons and the formation of glial scars around the electrodes, both of which degrade the recording signal-to-noise ratio (SNR) and stimulation efficacy^7–11^.

Flexible neural probes have emerged as a promising solution to these challenges by minimizing the mechanical mismatch between the probe and brain tissue^12–14^. We have previously demonstrated in NanoElectronic Thread (NETs) that when the thickness of polymer-based electrodes is reduced to about 1 µm, they become ultraflexible, enabling seamless electrode-tissue integration, stable month-long recordings, tracking of the same neurons^12,15^, and reliable chronic intracortical microstimulations^7^. However, compared with active silicon electrodes, flexible electrodes face a standing challenge in scalability, lacking the means to integrate a large count of channels within a small probe form factor. The limitation arises primarily from the low resolution of the photolithographic microfabrication process commonly used to define the feature sizes and interconnect density in these electrodes. Consequently, typical flexible electrodes^16,17^ contain 1-2 orders of magnitude fewer channels than the state-of-the-art active silicon probes of comparable width^4,5,18–20^. The constraints on channel density result in relatively low total channel numbers or sparsely distributed contact sites, limiting their capacity to distinguish densely clustered neurons over extended regions. Although total channel count can be increased by implanting multiple probes in parallel^16,21^, this strategy does not overcome the fundamental limitation in recording-site density and inevitably increases surgical complexity, tissue injury, and associated risks. Similarly, more channels can be accommodated by increasing the probe width; however, the larger implant footprint can exacerbate—and often cause permanent—tissue damage^22–26^, thereby compromising recording quality and stimulation efficacy.

Overcoming this density limitation while preserving the biocompatibility and chronic stability of flexible probes requires substantially increasing interconnect density without enlarging the implant. Here we report nanofabrication-enabled high-density NETs, in which drastically reduced interconnect linewidth and pitch enable a tenfold increase in channel counts while maintaining the few-micron thickness for ultraflexibility, and a shank width of 70 – 100 µm comparable to previously reported flexible probes ^16,24,27^ and rigid probes ^4,5,18–20^. To drastically increase the channel density of flexible electrodes, we systematically explored the design parameter space and developed an electron-beam lithography (EBL) strategy capable of patterning interconnect traces as narrow as 150 nm. This approach created a family of high-density polymer neural electrodes containing 128 to 512 channels while retaining the slender, ultrathin form factors of NETs^7,12,21,22,28^. We demonstrate that the resulting EBL-fabricated NETs (EBL-NETs) enable spatially oversampled recording, which is critical for precise isolation of densely clustered neuron populations while offering extended brain region coverage. Their recording stability permits reliable tracking of neuronal populations over 10 days without drift corrections. EBL-NETs further support spatially selective neural modulation together with simultaneous, high-resolution recording of evoked neuronal activity and functional connectivity. We establish the versatility of EBL-NETs across diverse electrophysiological applications, including longitudinal behavioral studies in mice and rats. Extending these capabilities toward human applications, we demonstrate stable intraoperative recording from human temporal cortex and identify lexical-semantic encoding in population activity. Together, these results establish EBL-NETs as a scalable, high-density, and chronically stable neural interface that bridges fundamental circuit studies in animal models with translational applications in the human brain.

## Results

### Design principles of EBL-NETs

We developed a multilayer lithography process to pattern high-density electrode arrays by electron-beam lithography (EBL). We define hundred-nanometer-wide metal interconnect traces by EBL and distribute them across two metal layers (Figure 1A). The structural backbone of the electrode incorporates four polyimide layers: a bottom encapsulation layer, a separation layer that enables two stacked interconnect layers, a top encapsulation layer for electrical insulation, and an electrode encapsulation layer that reinforces the electrode contact sites and prevents delamination during extended microstimulation sessions. This overall design balances the high channel density, nanofabrication complexity, and total device thickness, allowing EBL-NETs to scale to hundreds of channels within a thin, thread-like probe of minimal cross-sectional area.

**Figure 1.**
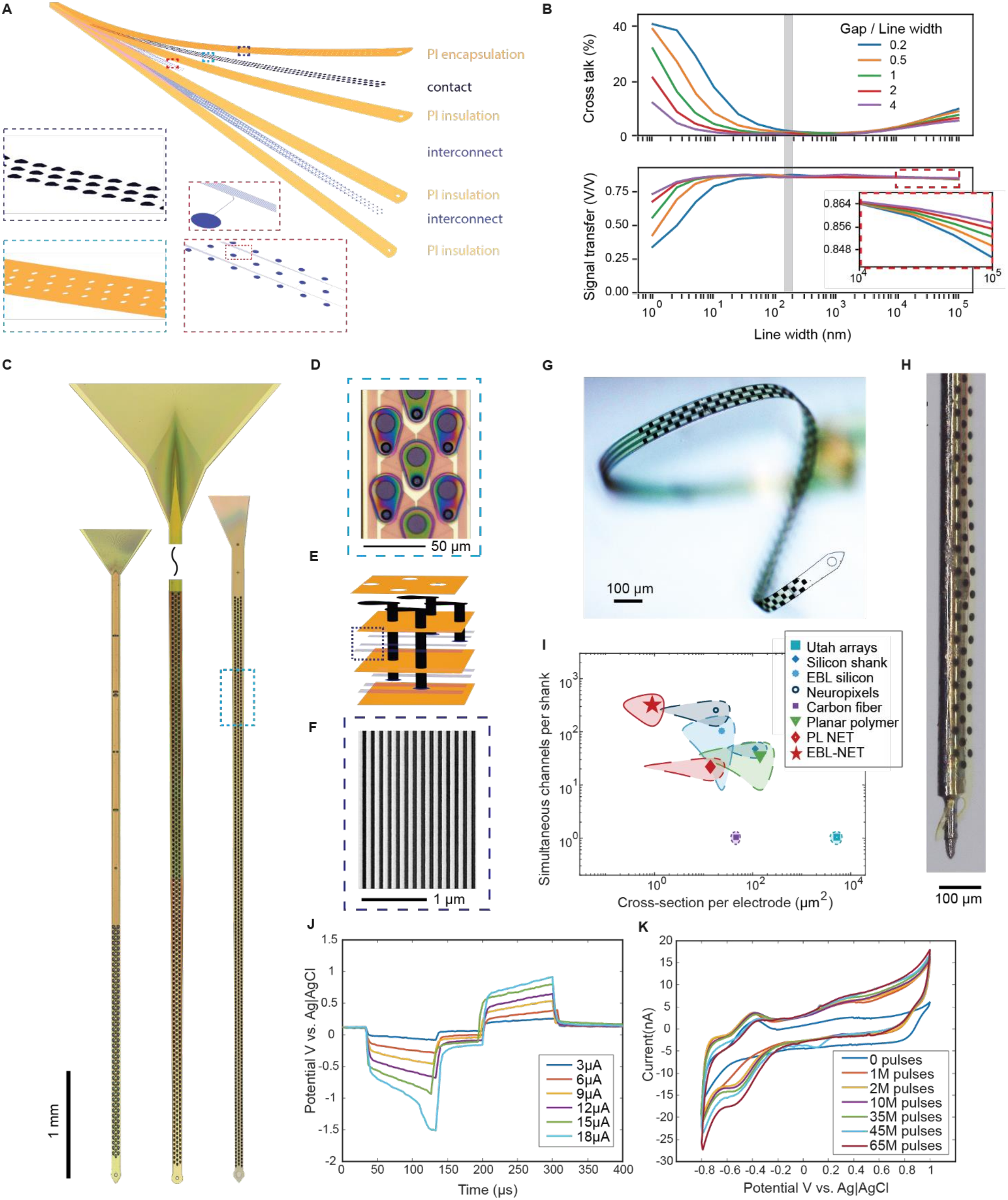
The design principle, fabrication, and characterization of ultraflexible EBL-NETs. (A) Structural schematic of EBL-NET, comprising three polyimide insulation layers, two interconnect layers, a contact layer, and an encapsulation layer. Insets highlight design details of individual layers. (B) Simulated crosstalk and signal-transfer as a function of linewidth and gap-to-linewidth ratios. The gray shaded region marks the 150–200 nm linewidth range that provides acceptable crosstalk and signal transfer. (C) Three representative designs: 128 channels with 2 mm contact coverage and 5–7 mm implantable length (left); 384 channels with 5 mm contact coverage and 10 mm implantable length (middle); and 512 channels with 5 mm contact coverage and 5.5 mm implantable length (right). (D) Zoom-in view of an EBL-NET showing circular contacts of 15 µm in diameter and via of 5 µm in diameter. (E) Schematic diagram showing that interconnect traces (gray) individually address contacts through vertical vias (black) and are insulated by polyimide encapsulation (orange). (F) Scanning electron microscopy (SEM) image of the interconnect traces of 150-nm linewidth and 150-nm gap. (G) An EBL-NET suspended in water, highlighting its ultraflexibility, high-density contacts, and a through hole (35 µm in diameter) at the distal end to facilitate implantation. (H) A 128-channel EBL-NET threaded on a sharpened tungsten wire for implantation. (I) Comparison of EBL-NETs with representative existing neural electrodes of various types in terms of their implantation footprints and scalability. Data points include Utah arrays^29,30^, silicon probes^31–33^, EBL silicon probes^34,35^, Neuropixels^4–6,36,37^, carbon fibers^38^, traditional polymer probes^39–41^, ultraflexible NETs fabricated via photolithography^7,21,22^, and EBL-NET (this work). (J) Representative voltage transients of biphasic pulses at varying current levels (3–18 µA). (K) Cyclic voltammetry (CV) after millions of 10 µA biphasic pulses.

Reducing linewidth and pitch enables higher channel counts to fit within a fixed shank width, but excessively narrow traces increase resistance and may compromise signal transfer. To predict EBL-NET performance and guide device design, we conducted simulations combining finite element and analytical circuit models (Figure S1). The resistance of the nanoscale traces could contribute tens of kiloohms depending on the width and length^42^ (Figure S1A), increasing the total series impedance between the electrode–electrolyte interface and the amplifier input and potentially attenuating recorded signals through voltage division arising from the amplifier input impedance. We defined an acceptable trace resistance as <10% of the total impedance at 1 kHz, corresponding to a 15 kΩ cutoff given the typical ∼ 150 kΩ impedance of our IrOx-coated contacts at 1kHz^43,44^. The minimum linewidth ranges from 80 – 320 nm for probe lengths of 5 – 20 mm (Figure S1A). We further modeled crosstalk and signal attenuation by accounting for capacitive coupling between adjacent traces, amplifier input impedance, traceline resistance, shunt impedance between the probe and surrounding tissue, and electrode/electrolyte interface^43^. Coupling and shunt parameters were extracted using finite-element modeling of both in-plane and out-of-plane neighboring traces and incorporated into a distributed two-line circuit model to simulate signal propagation and crosstalk (Figures S1B–S1D). Across all design-relevant geometries, both in-plane (Figure 1B) and out-of-plane (Figures S1E and S1F) neighboring traces exhibited low predicted crosstalk (<1%), and the signal transfer closely matched that expected from the voltage-divider effect due to the traceline resistance. Guided by these simulation results and balancing the tradeoff between interconnect density, crosstalk, and signal attenuation, we selected linewidth of 150 – 200 nm and an interline spacing of 150 - 250 nm for most electrodes used in this study as a balance between minimal footprint and fabrication process robustness. Specifically, 180-nm-wide, 10-mm-long traces with 220-nm spacing yield a predicted crosstalk of approximately 0.8% and signal attenuation of 13.4%.

Beyond electrical performance, linewidth selection also involves practical considerations in EBL fabrication. We therefore established writing-current guidelines that balance patterning resolution and fabrication throughput across different linewidths. E-beam patterns were divided into polygons, with exposure doses calculated based on proximity effects, clear dose, and dose coefficients. Higher writing currents reduce exposure time but increase pixel size, potentially compromising edge definition for features approaching the pixel dimensions. The resulting writing times, including overhead from stage movement, data transfer, and exposure were calculated (Figure S2A), from which we chose a 10 nA write current for a nominal linewidth of 150–200 nm, taking less than 50 s per 100 10-mm long traces.

We also considered whether nanoscale interconnects could withstand prolonged electrical stimulation. Mean time to failure (MTTF) was estimated under stimulation conditions representative of in vivo applications using a customized model informed by prior literature^44^(Figure S2B). Simulations used 100 Hz biphasic stimulation at 1–20 µA and predicted MTTFs exceeding 1,000 days for linewidths above 200 nm, indicating that interconnect failure is unlikely under typical stimulation conditions. Together with the electrical and fabrication constraints above, these analyses define a practical design space for EBL-NET interconnect geometry.

Informed by these design considerations and leveraging the maskless nature of EBL, we readily fabricate diverse designs with contact arrangements tailored to various brain regions and animal species. Figure 1C shows three representative EBL-NET designs, consisting of 128-, 384- and 512-channel contact arrays with depth coverages ranging from 2 to 5 mm. Each design includes a 35 µm hole at the tip for engagement with an insertion shuttle and contains three columns of densely patterned contacts with 64 – 102 channels per millimeter of depth, enabling spatially oversampled recordings that improve single neuron isolation^34,36^ (Figure 1C). Each contact is individually connected to an interconnect trace through a via in the polyimide encapsulation (Figures 1D and 1E).

EBL enabled patterning of metal interconnects only one or two hundreds of nanometers wide over the probe length, yielding a high aspect ratio of approximately 1:50,000 (Figure 1F). To improve fabrication yield at this extreme aspect ratio, we implemented double-pass EBL lithography using advanced alignment methods, achieving precise pattern alignment and clean lift-off even at angled turns (Figure S3). These methods resulted in an overall channel yield of 91.4% across 100 tested devices (Figure S4A). The resulting EBL-NETs have overall thicknesses ranging 2 – 5 µm, rendering them ultraflexible and unable to remain freestanding without support in either air or solution (Figure 1G).

Because EBL-NETs are too flexible to penetrate brain tissue independently, we incorporated a shuttle-assisted implantation strategy in which a micro-machined stiffener delivers the probe while minimizing the surgical footprint (Figure 1H). Compared with other flexible neural probes, EBL-NETs possess significantly increased number of simultaneously recording channels, approaching that of high-density silicon probes^4,5,38^ while retaining the small cross-sectional footprint of ultraflexible polymer probes^12,20,21^ (Figure 1I).

Lastly, we evaluated the robustness of charge injection through sputtered IrOx contacts. Biphasic current pulses from 3 to 18 µA produced current-dependent voltage transients, with currents up to 15 µA remaining within the water window^45^ (Figure 1J). Cyclic voltammetry (Figure 1K) and impedance spectroscopy (Figure S4B) remained stable over 65 million pulses of 10 µA biphasic stimulation, supporting the electrochemical robustness of the electrode interface for chronic intracortical microstimulation (ICMS).

### EBL-NETs enable high-density and stable chronic electrophysiological recordings

We primarily used mouse and rat models to validate the recording performance and chronic stability of EBL-NETs. Figure 2A presents a representative recording session from a rat, with a 128-ch EBL-NET implanted across the secondary motor cortex (M2) and cingulate cortex area 1 (Cg1) (Table S1). In a representative recording session (Figure 2A), high-pass filtered traces showed robust multi-channel spiking activity, with estimated single-unit positions distributed along the probe. We detected 112 units, with peak amplitudes of 77.2 +/- 31.9 µV and SNR of 11.92 +/- 4.93 (mean +/- SD, methods). The high contact density spatially oversampled individual units, with their extracellular waveforms detected across multiple neighboring contacts, as demonstrated by the representative waveform maps from two probe regions (Figure 2A).

**Figure 2.**
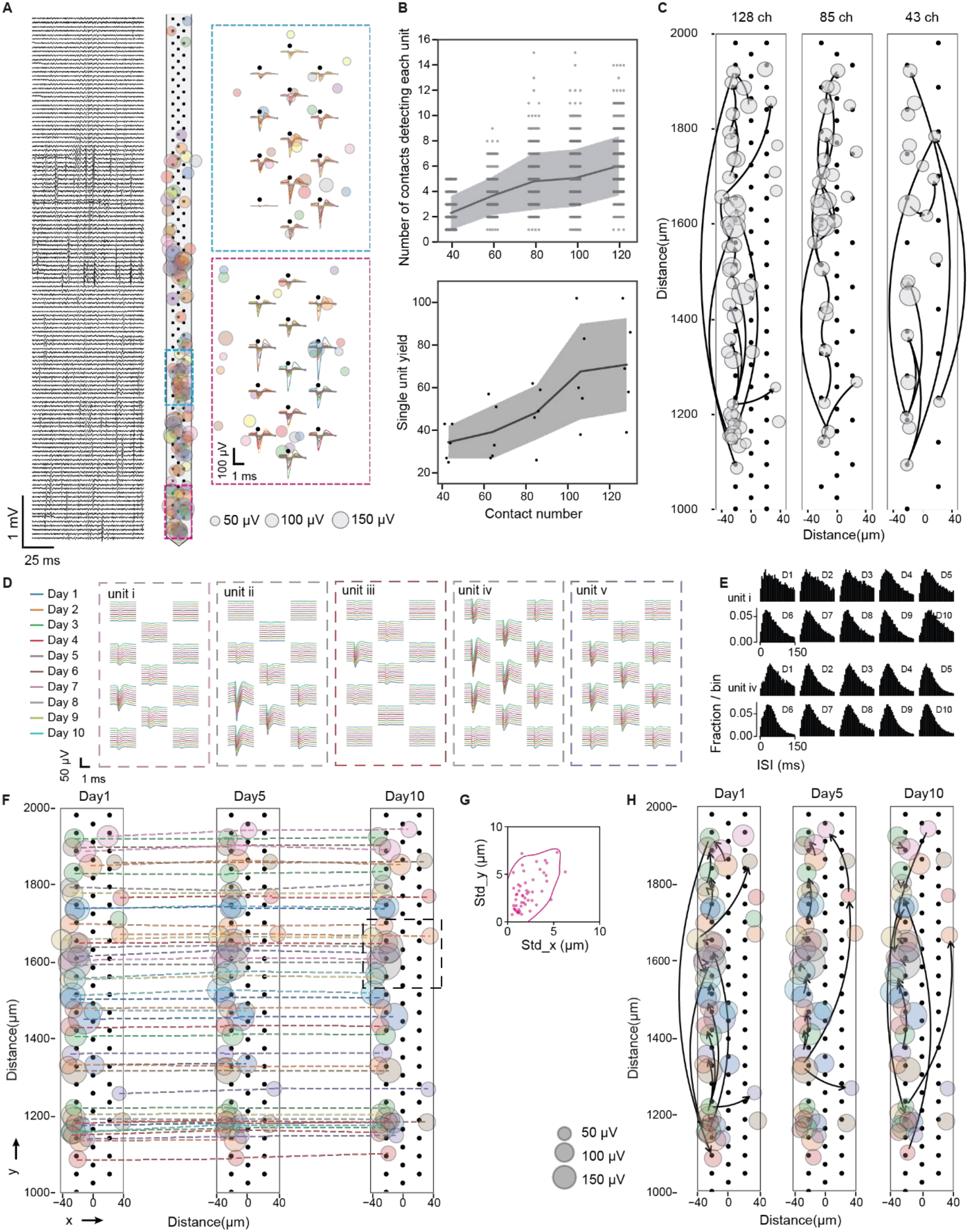
High-density EBL-NETs enhance neural population recording, single-unit and putative monosynaptic connectivity detection, and longitudinal tracking. (A) Representative extracellular recording from a 128-ch EBL-NET. Left, high-pass filtered recording signals. Middle, estimated locations of single units overlaid on the contacts (black dots). Right, examples of waveform distributions from two representative sections, depicting mean spike waveforms across nearby contacts and the corresponding unit-location (filled circles). Color presents units; circle size indicates waveform amplitude. (B) Contact density enhances single-unit detection. Top, number of contacts detecting each unit increases with contact density; each dot represents one unit. Bottom, single-unit yield as a function of contact number. Each dot represents one recording session; line and shading show mean and standard deviation. (C) Impact of contact density on estimating unit location and detection of putative monosynaptic connectivity. Gray circles indicate unit location, black dots indicate contacts, and directed curves indicate putative monosynaptic connections between unit pairs (Methods). (D) Representative longitudinal waveform stability for five tracked units over 10 days. Each subpanel shows mean waveforms from day 1 through day 10, with days indicated by line color. (E) Inter-spike interval distributions for two tracked units in (D). (F) Estimated locations of 55 trackable units (Shaded circles with diameter indicating peak amplitude) relative to the NET across days 1, 5, and 10 from the same recording in (D) and (E), with dashed lines connecting the same units to depict minimal position drift. (G) Longitudinal location drift of the 55 tracked units measured as the standard deviation (SD) of the unit’s estimated locations across 10 days. (H) Putative excitatory monosynaptic connectivity maps on day 1, day 5, and day 10. Directed curves indicate curated source-target pairs from CCG-based connection analysis.

To quantify how electrode density influenced recording performance, we synthesized datasets from 128-ch EBL-NETs recordings (n=5 mice; Table S1) using evenly reduced contact layouts containing 43, 64, 85, and 106 channels and compared their recording outcomes with the original 128-ch layout. Subsampling analysis confirmed the benefit of spatial oversampling: the number of contacts detecting each unit increased with contact density (Figure 2B), supporting more reliable single-unit detection and isolation. Consistently, single-unit yield increased from 34.4 +/- 8.5 accepted units at 43 contacts to 70.8 +/- 24.4 at 128 contacts (mean +/- SD, n = 5 mice). The estimated unit locations, determined by triangulation^46^, progressively shifted as contact density was reduced (Figure 2C) and fewer contacts sampled each neuron and provided less spatial information for localization. This loss of units and spatial information also affected downstream circuit inference. Putative excitatory monosynaptic connections identified from curated cross-correlogram (CCG) (Methods, Figure S5) decreased in number and exhibited altered spatial distribution with fewer contacts.

The high electrode density also enhances the fidelity of tracking large populations of neurons over multiple recording sessions. We examined the longitudinal stability of EBL-NETs using a representative 10-day recording series from the somatosensory cortex of a mouse. We sorted and tracked units that maintained similar multi-contact waveform patterns across days (Figure 2D). We verified the tracking results by confirming that the inter-spike interval (ISI) distributions from tracked units remained consistently structured over the same 10-day period (Figure 2E). A total of 55 units were tracked across days (Figure 2F), with highly stable inferred location. Across these tracked units, the median standard deviations in estimated locations were 1.75 μm laterally across the probe width and 2.91 μm vertically along the implantation depth, with 92.7% of units exhibiting standard deviations below 6 μm and 8 μm in these respective dimensions (Figure 2G). Recurring putative source-target connections were observed in the longitudinal recordings, alongside the emergence and disappearance of individual connections (Figure 2H). Over longer timescales of months, EBL maintained stable performance, including stable channel impedance and unit amplitudes. (Figure S6). Post-mortem histology four months after implantation showed tight tissue-probe integration, with preserved neuronal and nuclear distribution around the EBL-NET, no apparent elevation of microglia density, and minimal astrocytic responses (Figure S7). Collectively, these results demonstrate that high-density EBL-NETs enhance single-unit detection and spatial localization, improve the inference of local functional connectivity, and enable chronically stable monitoring of both unit activity and network connectivity.

### EBL-NETs provide high-resolution ICMS with spatially selective neuronal activation

Having established the high-density recording capability of EBL-NETs, we next leveraged the same IrOx-coated electrode array for simultaneous ICMS and recording to directly resolve stimulation-evoked neuronal and network responses. Because ICMS activates neurons primarily through axons^47^, its spatial selectivity depends strongly on the channel spacing of the electrode arrays. The high-density EBL-NETs may therefore enable fine-grained modulation of neuronal activity. To test this capability, we performed simultaneous ICMS and electrophysiological recording using 128-ch EBL-NETs implanted in mouse somatosensory cortices. In a representative session (Figure 3A), we selected ten channels along the probe for stimulation, delivered pulse trains at currents of 1 – 5 µA, and simultaneously recorded neural activity from all 128 channels. Although stimulation artifacts were apparent in the raw recordings (Figure 3B), stimulation-evoked spikes were readily detected and isolated by spike sorting after suppression of stimulation artifacts (Methods). Consistent with their seamless integration with neural tissue, EBL-NETs elicited neuronal activation at low currents of 2 µA in all animals tested (n=5, Figure S8), comparable to photolithographically fabricated, lower-density NETs^7^ and substantially lower than the currents typically required by rigid electrodes^7^.

**Figure 3.**
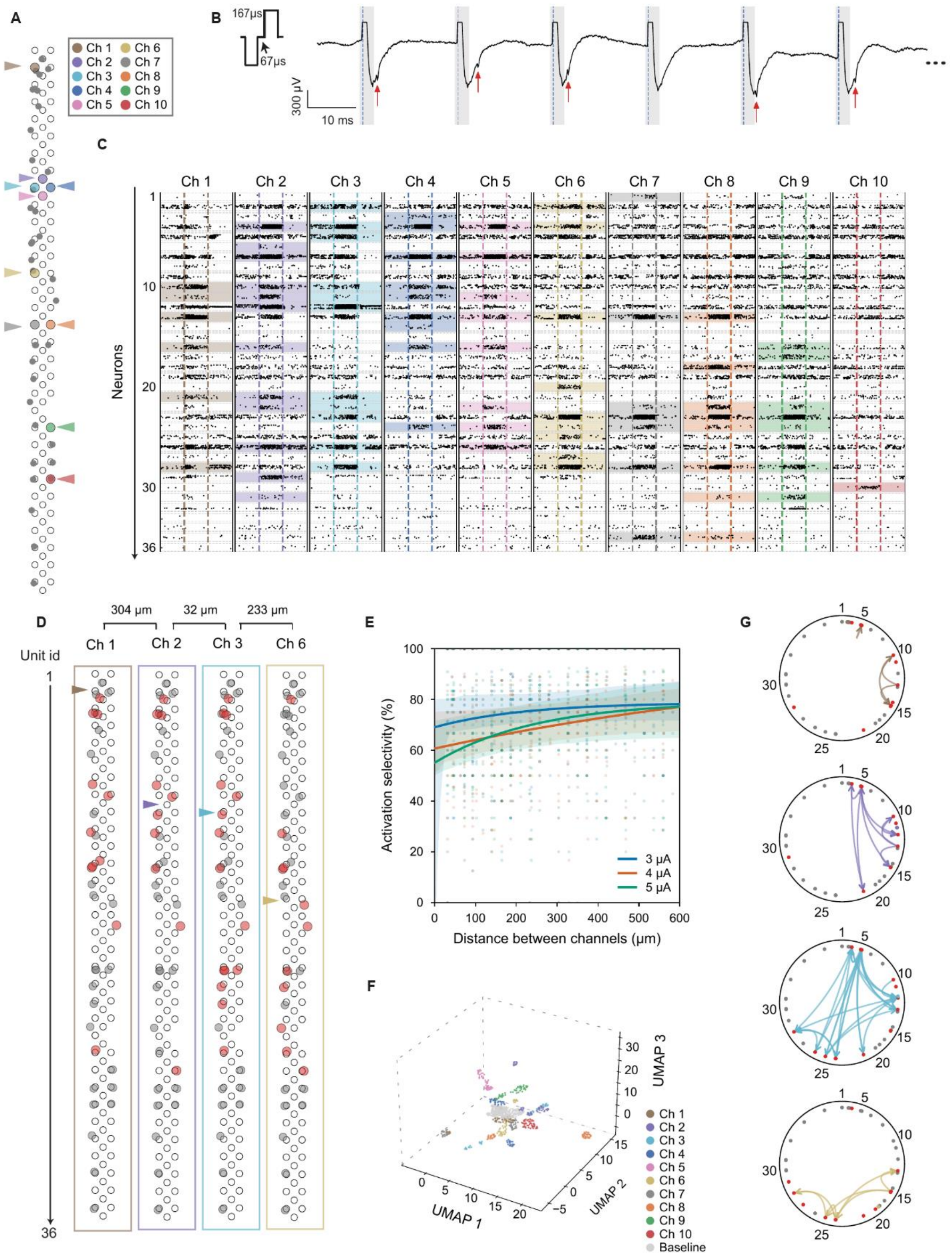
Bidirectional EBL-NETs enable spatially selective stimulation with simultaneous recording of neuronal and network responses. (A) Representative ICMS and simultaneous single-unit recording by EBL-NET, showing estimated unit locations (filled circles) along the probe and the ten channels that were used for stimulation. Colored arrows denote stimulation channels. (B) Representative raw recording trace during ICMS consisting of six 5 µA pulses (marked by vertical lines). Gray bands denote artifact-blanking windows immediately following the pulses; red arrows mark detected spikes from an isolated unit. (C) Spike rasters of 36 isolated units to 5 µA ICMS delivered through each of ten channels. Rows represent curated units ordered by estimated position along the probe, and each block shows ten ICMS trials for the indicated stimulation channel. Colored backgrounds mark units with significantly increased firing during stimulation relative to baseline. Dashed vertical lines mark the stimulation onset and offset. (D) Spatial distribution of activated units of four representative stimulation channels (marked by colored arrows). Each column shows position-ordered units; red and gray circles indicate activated and non-modulated units, respectively. The distance between the stimulation contacts are marked at the top of the columns. (E) Activation selectivity as a function of distance between stimulation channels. Activation selectivity was defined as 100% minus the percentage overlap of activated units between pairs of stimulation channels. Points show channel-pair measurements at 3, 4, and 5 µA; curves and shaded bands show per-current exponential fits with 95% bootstrap confidence intervals. (F) UMAP representation of population activity during stimulation (color coded by stimulation channel) and baseline (gray). (G) Stimulation-induced functional coupling for the same representative channels. Dots indicate position-ordered units arranged around the circle, with numeric labels indicating unit identity. Directed curves denote unit pairs with short-latency temporal relationships in stimulation-aligned cross-correlograms, suggesting functional coupling (Methods).

We next examined the spatial selectivity of ICMS from simultaneously recorded population responses. Different stimulation channels recruited overlapping but distinct subsets of neurons (Figure 3C). Figure 3D illustrates the spatial selectivity for four representative stimulation sites, including two neighboring channels separated by ∼30 µm center-to-center, and two more distant channels. Even the closest neighboring contacts evoked partially distinct activation patterns, with responses becoming increasingly distinct as the separation between stimulation sites increased. We quantified spatial selectivity as a function of stimulation contact distance using an activation-selectivity metric, defined as 100% minus the percentage overlap of activated units between two stimulation sites at the same current. Activation selectivity increased with contact separation and decreased with stimulation amplitude (n=5, Figure 3E). Notably, activation selectivity remained above 50% at all currents and above 70% at 3 µA, even at the closest stimulation-site separation.

We next examined population responses to stimulation. Uniform Manifold Approximation and Projection (UMAP) embedding of recordings during baseline and ICMS revealed distinct clusters associated with individual stimulation sites (Figure 3F), demonstrating separable population responses to stimulation at different sites. Stimulation-associated cross-correlogram (CCG) analysis (Methods) further identified stimulation-site-dependent patterns of putative short-latency functional coupling among stimulation-responsive units (Figure 3G). Despite a small fraction of overlapping connections, nearby stimulation sites engaged predominantly site-specific source units and putative functional connectivity (n=5 mice, Figure S9), supporting stimulation-site-specific network responses from high-density EBL-NETs.

We further examined the longitudinal stability of ICMS-evoked neuronal responses by conducting 7 stimulation sessions spanning two weeks, each with the same 10 stimulation channels as shown in Figure 3C. Twenty-one units were tracked across all the sessions, with 14 units having elevated firing rate during stimulation, and 51 significant stimulation channel -unit pairs. A representative tracked unit maintained a consistent spatial waveform profile spanning multiple contacts and a reproducible short-latency response to ICMS across sessions (Figure S10). Across tracked stimulation channel - unit pairs, activation thresholds spanned from 1.2 to 4.8 µA and remained stable over time, with a maximum SD 1.2 µA across sessions (Figure S11). The magnitude of stimulation-evoked modulation was similarly stable after normalization across sessions, with mean modulation index ranging from 0.590 to 0.924, and a SD of modulation index below 0.2 for all pairs (Figure S12).

Together, these results demonstrate that high-density EBL-NETs offer spatially selective ICMS through closely spaced contacts, simultaneous high-resolution measurement of neuronal and local-network responses, and stable neuronal recruitment across longitudinal sessions, providing a scalable platform for precise modulation and interrogation of local neural circuits.

### EBL-NETs provide stable chronic recordings in behaving rodents

We next evaluated EBL-NETs for high-density chronic recordings in longitudinal neuroscience studies spanning multiple months. In a rat action sequence task, animals were trained to perform a left-left-right (L-L-R) sequence of side-port entries for reward while an EBL-NET recorded from the anterior cingulate cortex (ACC). In a representative session, 104 simultaneously recorded units exhibited reproducible sequential activity associated with repeated left and right actions, visualized by Rastermap^48^-ordered (Methods) population activity (Figure 4B and Figure S13). Action-related activity persisted longitudinally: a representative choice-modulated neuron maintained selective responses to right actions from post-implantation day 115 to day 180 (Figure 4C). Across this chronic period, 23 single units were longitudinally tracked (STAR Methods), with stable left-versus-right decoding scores through all sessions (Figure 4D). At the population level, projection of tracked unit activity from all sessions into day-115 principal-component space preserved representations of the first left, second left, and right sequence steps (Figure 4E). On top of that, we trained a three-class linear discriminant analysis (LDA) classifier using day-115 PC1 and PC2 scores to classify the first L, second L, and R sequence steps and evaluate the stability of population geometry in later sessions (STAR Methods). The same decision boundaries shifted by session-wise population mean yielded classification accuracies of 0.98, 0.99, and 0.91 in days 116, 120, and 180, respectively. These results support the preservation of action-related single-unit selectivity and ACC ensemble geometry across months, consistent with previous observations of long-term stability in behavior-related single-neuron activity and stable population-level representations^49–51^.

**Figure 4.**
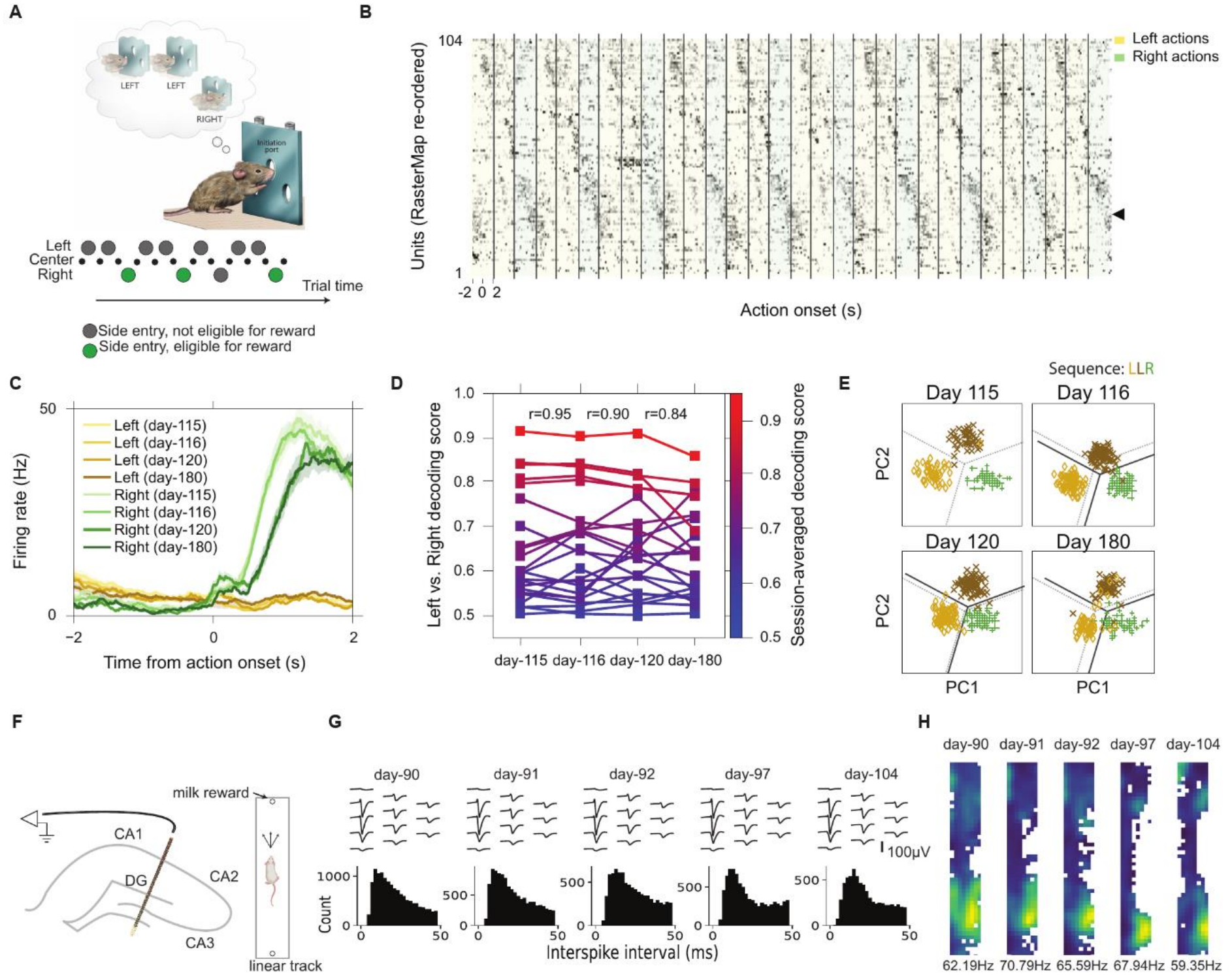
Longitudinal tracking of single-unit and population activity in behaving rodents. (A) Schematic of the rat behavioral task to study action sequence encoding in anterior cingulate cortex (ACC). (B) Sequential ACC population activity during 30 consecutive actions. Rastermap^48^-ordered activity of 104 simultaneously recorded units aligned to action onset. Background shading denotes left (yellow) and right (green) actions. (C) Action-aligned mean firing rate of a representative choice-modulated, longitudinally tracked ACC unit during left (yellow) and right (green) actions on post-implantation days 115, 116, 120, and 180. Shading denotes the standard error of the mean (s.e.m.). (D) Left-versus-right decoding scores of 23 longitudinally tracked single units. Lines connect individual units; color indicates session-averaged decoding score. Adjacent-session Pearson correlations were 0.950, 0.904, and 0.843. (E) Principal-component representations of population activity associated with the first left, second left, and right actions across sessions. Activity from all sessions was projected onto the first two principal components from day 115, and a three-class linear discriminant analysis decoder trained on day 115 was applied to later sessions. Dotted and solid lines show the day-115 decision boundaries and the boundaries translated by each session’s population-mean shift, respectively. (F) Schematics of EBL-NET implantation and the tethered linear-track navigation task (right). (G) Mean waveforms and ISI distributions of a representative, longitudinally tracked CA1 place cell on post-implantation days 90, 91, 92, 97, and 104. (H) Spatial firing-rate maps for the same unit across sessions. Peak rates are shown below each map; white bins indicate unvisited or masked spatial bins.

We further evaluated the longitudinal recording stability of place cells in the rat hippocampus. An EBL-NET spanning the posterior parietal cortex, CA1, and dentate gyrus recorded neural activity during tethered linear-track spatial alternation task (Figure 4F). A representative hippocampal place cell was tracked across post-implantation days 90, 91, 92, 97, and 104 with stable waveforms, consistent ISI distributions, and similar spatial firing fields (Figures 4G and 4H). In a separate mouse reach-and-grasp study, an EBL-NET implanted in the primary motor cortex similarly recorded stable task-related activity across consecutive training days (Figure S14). Together, these studies demonstrate longitudinal tracking of single neurons and population activity across multiple brain regions, behavioral paradigms, and rodent species.

### High-density intraoperative single-unit recording in human temporal cortex

Finally, we evaluated the translation feasibility of EBL-NETs for human neuroscience. Cardiac pulsation and respiration present a significant challenge for maintaining a stable probe-tissue interface in human cortex. Substantial probe–tissue motion comes from the mechanical mismatch between rigid probes and brain tissue, compromising recording stability and device integrity^52,53^. To facilitate intraoperative deployment, we designed a ready-to-use package housing the EBL-NET and bonded recording amplifier within a sterilizable metal enclosure (Figure 5A). We positioned the enclosure with an engagement pole away from the penetrating tip, leaving the shuttle wire unobstructed for retraction (Figure 5A inset). In a representative recording session, two EBL-NET probes were simultaneously inserted to a depth of ∼4 mm in a portion of the left anterior middle temporal gyrus that was planned for resection during a left anterior temporal resection in a patient with medically refractory epilepsy (Figure 5B). High-density recordings resolved large numbers of neurons distributed along the probe (109 and 100 units on the two EBL-NET after shuttle wire retraction, respectively), with individual extracellular waveforms sampled across multiple neighboring contacts (Figure 5C).

**Figure 5.**
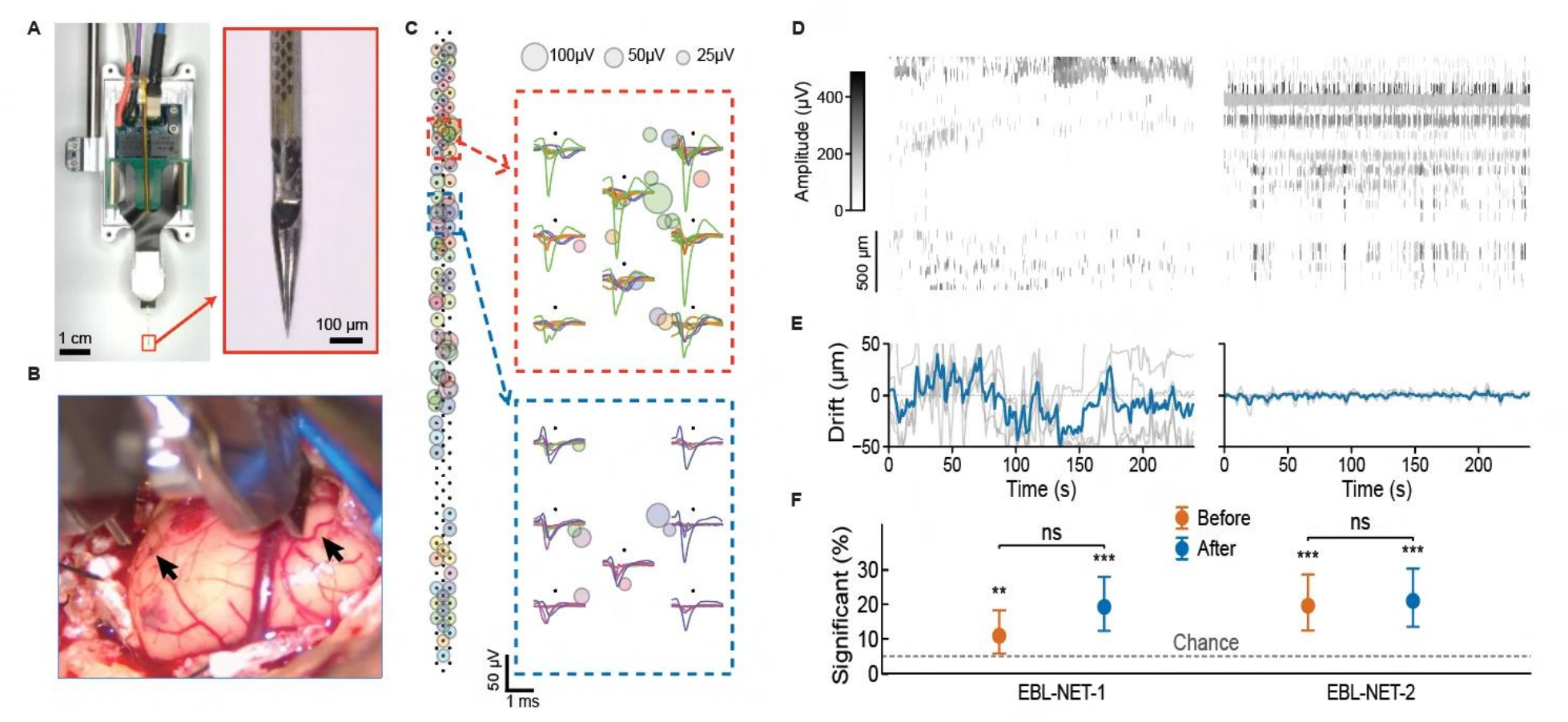
High-density single-unit recording in the human brain. (A) Photo of a ready-to-deploy EBL-NET package. Inset, NET engaged with a micro-machined tungsten shuttle wire. (B) Two EBL-NETs (pointed by the black arrows) implanted intraoperatively into human temporal cortex. (C) 100 single units recorded by EBL-NET-2 after shuttle wire retraction, with estimated locations overlaid on the contact layout. Circle area is proportional to peak-to-peak waveform amplitude; black dots indicate contacts. Dashed callouts show representative local waveform groups and their associated units, indicated by matching colors. (D) DREDge-localized threshold-crossing events during 240-s recording sessions before (left) and after (right) shuttle-wire retraction, plotted by inferred depth, with grayscale intensity indicating absolute peak amplitude. (E) DREDge estimates of probe displacement before (left) and after (right) shuttle-wire retraction. Gray curves show four local nonrigid DREDge displacement estimates along probe depth; blue shows their mean. (F) Percentage of units recorded by the two EBL-NETs showing significant lexical-semantic encoding before and after shuttle retraction. Points show count-derived percentages; error bars show 95% Clopper-Pearson confidence intervals. All four conditions exceeded the 5% chance level (one-sided exact binomial tests; EBL-NET-1: before, 12/110, 10.91%, P = 0.009; after, 21/109, 19.27%, P < 0.001; EBL-NET-2: before, 20/102, 19.61%, P < 0.001; after, 21/100, 21.00%, P < 0.001). Before-versus-after differences were not significant for either probe (two-sided Fisher’s exact tests; EBL-NET-1, P = 0.092; EBL-NET-2, P = 0.862).

We next examined whether the ultraflexible structure of EBL-NETs reduced relative probe–tissue motion by contrasting recordings before and after shuttle-wire retraction. DREDge-localization of threshold-crossing events^54^ (Methods) revealed substantially reduced displacement after shuttle retraction (Figure 5D). The maximum absolute mean probe drift decreased fivefold, from 48.9 µm to 8.9 µm (Figure 5E), consistent with improved mechanical coupling between the ultraflexible probe and the moving human brain after removal of the rigid shuttle.

We analyzed lexical-semantic encoding during passive listening to podcasts under general anesthesia using the large population of single units recorded by EBL-NETs. Regularized Poisson generalized linear models predicted word-evoked spike counts from Word2Vec-derived features and word duration^55,56^ (Methods). For both probes, the percentage of neurons showing significant lexical-semantic encoding was significantly higher than chance level both before and after shuttle retraction (Figure 5F). Notably, the total number of neurons exhibiting significant lexical-semantic encoding increased from 32 to 42 after shuttle-wire retraction, confirming that EBL-NETs retained access to functionally informative single-unit activity without rigid insertion support. Together with the substantially reduced relative probe–tissue motion, these results support the feasibility of high-density, functionally informative neural recording in the human brain using ultraflexible interfaces.

## Discussion

The central advance of EBL-NETs is the integration of high channel counts and dense spatial sampling within an ultrathin polymer architecture that supports stable chronic recording and electrical stimulation. Whereas previous flexible neural interfaces established the advantages of minimizing probe dimensions and mechanical stiffness^21,24,28^, our wafer-scale EBL fabrication substantially increases channel count and spatial sampling density while retaining the ultraflexible form factor. By combining nanoscale interconnects with multilayer routing, we fabricated probes of 128–512 individually addressable contacts. Dense spatial sampling improved detection and isolation of closely spaced single units, while the mechanical compliance of EBL-NETs enabled stable longitudinal tracking of neuronal populations. The same high-density architecture further supported spatially selective ICMS with simultaneous measurement of neuronal and network responses. These capabilities were demonstrated across longitudinal studies in behaving mice and rats, and extended to high-density single-unit recording in the human brain. Together, EBL-NETs substantially expand the spatial sampling and perturbation capabilities of flexible neural interfaces, providing a scalable platform for high-resolution interrogation of neuronal populations across experimental models and the human brain.

High-density silicon probes such as Neuropixels have substantially advanced neural population recording through integrated electronics and dense spatial sampling^5,36^. However, their rigid silicon shanks remain mechanically mismatched to soft, continuously moving brain tissue, a limitation that increased recording-site density alone cannot resolve. Relative probe-tissue motion can alter extracellular waveforms or lead to signal loss, complicating reliable identification and longitudinal tracking of individual neurons. Mechanical mismatch can generate interfacial strain and contribute to chronic neuroinflammation and neuronal loss around rigid implants^54,57^. These challenges become particularly consequential in the human brain, where substantial physiological motion can compromise recording stability and necessitate motion compensation^54,58^. In contrast, the ultrathin polymer architecture of EBL-NETs is designed to accommodate tissue motion and reduce interfacial forces. EBL-NETs therefore integrate two complementary requirements for high-resolution neural interfaces—dense access to neuronal populations and mechanical compatibility with neural tissue, offering potential advantages in large brains and for chronic implantations.

Our results demonstrate the value of combining recording density with stability. Increasing the number of available contacts increased unit yield and provided richer multichannel waveform signatures for unit localization and longitudinal matching. Because these comparisons were performed using subsets of the same recordings, they isolated the information gained from denser spatial sampling from differences in tissue response or recording conditions between separately implanted probes. The contact layouts examined here provide a useful balance between local sampling density and depth coverage, although they do not establish a universally optimal contact pitch. Importantly, the benefit of dense sampling extends over time: stable recordings allowed individual neurons and population activity to be tracked over weeks to months and related to behavior. EBL-NETs thus combine spatially resolved access to neuronal populations with longitudinal stability needed to examine circuit dynamics at both the single-unit and population levels.

Dense, individually addressable contacts also enable spatially resolved electrical manipulation of neuronal populations. Previous work established low-threshold, chronically stable stimulation with ultraflexible electrodes^7^; EBL-NETs extend this capability by combining closely spaced stimulation sites with simultaneous high-density population recordings. Stimulation through contacts separated by 30 µm recruited partially distinct ensembles, with activation patterns becoming increasingly distinct as stimulating-site separation increased. Stimulation-associated functional coupling similarly depended on stimulation site, indicating that spatially localized perturbations can engage distinct patterns of local network activity. However, because the first milliseconds after stimulation pulses were excluded from analysis, these measurements may not capture the earliest directly evoked spikes. Therefore, selectivity was characterized within the sampled population and analyzed response window rather than the complete set of directly activated neurons. More broadly, combining dense recording and spatially resolved stimulation provides a means to examine how local perturbations propagate through neuronal circuits. Repeated access to the same neuronal population for both recording and manipulation may further enable longitudinal investigations of the stability and plasticity of circuit responses^59^.

The combination of ultraflexibility and high-density sampling is particularly compelling for extending high-density single-unit recording to the human brain, where physiological motion presents a substantial challenge for penetrating probes. The marked reduction in probe-tissue drift following shuttle-wire retraction is consistent with the intended mechanical advantage of the ultraflexible architecture and its ability to accommodate brain motion during intraoperative recording. Beyond recording stability, the large number of simultaneously isolated single units enabled analysis of lexical-semantic encoding at the neuronal population level, extending prior single-cell studies of semantic representations in the human brain^55,56^. These results support the feasibility of using EBL-NETs to investigate human neural computations with single-unit resolution and dense population sampling. Together with stable chronic recordings in animal models, the demonstration of large-scale intraoperative single-unit recording in humans provides a strong foundation for extending EBL-NETs to longer-term human applications, including large-scale single-unit recording during prolonged monitoring in epilepsy units and chronic bidirectional brain–computer interfaces. Such applications could enable longitudinal investigations of human neural computations and provide high-resolution neural readout and spatially selective stimulation for restoring lost neurological functions.

Collectively, the present results establish EBL-NETs as a platform for dense neural recording and spatially resolved microstimulation, enabling longitudinal interrogation of neural circuits and providing a foundation for future high-bandwidth brain interfaces. At the current scale, fabrication throughput, device yield, interconnect reliability, and packaging are compatible with research use and small-scale production, providing a practical foundation for broader deployment. Further scaling of recording and stimulation capabilities, however, will require coordinated development of the probes and their supporting electronics. Passive nanoscale routing preserves the ultrathin penetrating structure, yet it still requires external connections, acquisition channels, and stimulation sources that ultimately constrain system-level scalability. Hybrid polymer-chip architecture represents a promising path toward integrating high-density electronics while retaining a mechanically compliant tissue interface^60^.

## Resource availability Lead contact

Requests for further information and resources should be directed to and will be fulfilled by the lead contact, Lan Luan.

## Materials availability

This study did not generate new animal lines or unique reagents.

## Data and code availability

Figure-reproduction scripts, selected source and processed data used to generate the figures, and reference plots are publicly available in the EBL manuscript reproduction repository (https://github.com/Wangweinan16/EBL-manuscript-reproduction).

## Acknowledgments

We thank Nitin Tandon (McGovern Medical School, UTHealth Houston) for his support and contributions to the human intraoperative recording studies. We also thank Thomas Hamre, Sarah Soubra, and Nisha Giridharan for assistance with experimental setup, session logistics, and probe implantation, respectively. We thank Hannah Guffrey for help developing a pipeline to quantify fluorescence intensity and cell counts in histological sections. This work was supported by the National Institutes of Health (R01NS102917, U01NS115588, U01NS131086, R01EY036094, and U01NS121472), the Howard Hughes Medical Institute, the McNair Foundation, and the Gordon and Mary Cain Pediatric Neurology Research Foundation.

## Author contributions

Conceptualization: C.X. and L.L.; Methodology—electrode fabrication: W.W., Y.M., P.Z. and L.S.; implantation: Y.M., P.Z., L.S., H.W., K.H.L., R.Y. and X.Z.; Investigation and formal analysis—electrical modelling and reliability: W.W.; device characterization: Y.M. and P.Z.; mouse recordings: Y.M., W.W. and L.S.; microstimulation: Y.M., W.W., R.K., J.Z. and Y.L.; histology: L.D.L.P.; mouse reaching studies: R.Y. and Y.S.; rat frontal-cortex studies: H.W.; rat hippocampal studies: K.H.L. and P.A.; clinical coordination and human recordings: P.Z., K.A.K., C.A.K., H.Z., N.P., G.R. and S.A.S.; human-data analysis: H.Z. and E.A.M.; Visualization: Y.M. and W.W., with input from C.X. and L.L.; Supervision— device development: C.X. and L.L.; animal studies: A.Y.K., L.M.F., L.L. and C.X.; clinical studies and human-data analyses: S.A.S. and B.Y.H.; Writing—original draft: Y.M., W.W., L.S., L.L. and C.X.; Writing—review and editing: L.L. and W.W., with input from all authors.

## Supplemental information

The accompanying supplementary information contains Figures S1–S14 and Table S1.

## Methods

### Experimental model and study participant details

Mouse neural-recording experiments were approved by the Rice University Institutional Animal Care and Use Committee under protocol IACUC-22-200. Mouse intracortical microstimulation experiments were approved under protocol IACUC-22-202. All procedures were performed in accordance with the approved protocols and applicable institutional guidelines.

Two rats were used to demonstrate the longitudinal recording capability of the neural probes during behavior. Animal cohorts are summarized in Table S1.

One patient undergoing left anterior temporal lobe resection for medically refractory epilepsy participated in the intraoperative study. The study was approved by the

Institutional Review Board for Baylor College of Medicine and Affiliated Hospitals, under protocol H-56197. The EBL-NETS recordings were determined by the FDA and our IRB to be a nonsignificant risk (NSR) device study (G240178). Written informed consent for research participation was obtained from the participant before surgery.

## Method details

### EBL probe design and fabrication

EBL-NETs were fabricated using a multilayer polyimide (PI) process combining conventional photolithography for microscale features with electron-beam lithography (EBL) for nanoscale interconnects. The 128-channel EBL-NET contained a single interconnect layer, whereas the 384- and 512-channel designs used two interconnect layers separated by a 200-nm PI insulation layer. This multilayer routing enabled the probe shank width to be maintained at 80–120 µm. The fabrication process was based on that described in our previous work^7,12^, with interconnect layers patterned by EBL instead of photolithography for improved feature resolution. The PI layer thicknesses were selected to produce a mechanically balanced stack and minimize the distance of the metal interconnects from the neutral mechanical axis, thereby reducing bending-induced strain in the metal traces. The fabrication sequence described below corresponds to a representative ∼2-µm-thick device containing two interconnect layers.

Device fabrication was performed on a silicon substrate with a sacrificial Ni layer for subsequent probe release. PI 2574 (HD MicroSystems) was spun onto the wafer, baked on a 90 °C hot plate for 2 min, and then fully cured at 380 °C in a convection oven overnight to form a 0.9-µm-thick bottom PI layer that provided structural support and electrical insulation for the metal interconnects.

Nanoscale interconnects connected the electrode contacts to the BGA pads, with linewidth varied along the probe from approximately 150 nm near the electrode array to 750 nm near the BGA, where routing density was lower. The traces were widened progressively in regions with lower routing density to improve electrical conductance and fabrication robustness while retaining nanoscale dimensions where high routing density was required. The interconnects were patterned by EBL using an Elionix G-100 system operated at 100 kV, with a double-pass procedure. Each EBL pass was performed as follows. An MMA-based lift-off resist (AR-P 617.03) was first applied, followed by the positive-tone EBL resist CSAR 62 (AR-P 6200.09). A conductive polymer layer (Electra 92, AR-PC 5090) was coated over the resist stack to reduce charging during electron-beam exposure. Interconnect patterns were proximity-effect corrected using BEAMFOX with an initial clearance dose setting of 250 µC/cm^2^, followed by dose calibration in WECAS. Writing fields of 100–1000 µm were selected according to the alignment requirements of individual device regions. A beam current of 10 nA was used for features smaller than 250 nm, whereas 50 nA was used for features larger than 250 nm. Following exposure, the wafer was rinsed in deionized water for 60 s to remove the conductive layer, developed sequentially in AR 600-546 for 60 s and AR 600-50 for 20 s to develop the two resist layers, and then immersed in AR 600-60 stopper. The wafer was then blown dry and plasma cleaned using a Plasma Etch PE-10 before deposition of a 5-nm Ti adhesion layer and a 50-nm Au interconnect layer using a Sharon electron-beam evaporator. Lift-off was subsequently performed to define the nanoscale metal traces. All metal-and PI-deposition steps were preceded by plasma cleaning to remove resist residues and activate the underlying surface.

A double-pass EBL process was implemented to reduce discontinuities in the nanoscale interconnects. After the first interconnect pattern was defined using the EBL, metallization and lift-off sequence described above, a second EBL process was performed using an identical interconnect pattern aligned to the first pattern with the alignment functionality of the Elionix system. The second pass produced a 5-nm Ti/50-nm Au/5-nm Ti metal stack aligned with the first-pass traces. For devices containing two interconnect layers, the first double-pass interconnect layer was encapsulated with a 200-nm PI dielectric layer before fabrication of the second interconnect layer. The second layer contained a distinct routing pattern and was fabricated using the same double-pass EBL process.

The 500-µm-pitch ball-grid-array (BGA) pads were defined using an NR9-3000 (Futurrex, Inc.) resist mask and metallized with a Ti/Ni/Au stack for subsequent solder-ball bonding to the backend electronics. The interconnect layers and bonding pads were then encapsulated with a 0.7-µm PI layer. Via structures were defined with photoresist PR1-7000A1 (Futurrex, Inc.) and transferred into the PI layer by oxygen-plasma dry etching using a Plasma Etch PE-100, with the photoresist serving as an etch mask. The photoresist mask and etch residues were then removed with acetone before the electrode contacts were defined using an LOR/AZ resist stack. A Ti/Pt/Ti/IrOx electrode stack was deposited by sputtering, making electrical contact with the exposed landing pads of the interconnect layer through the PI vias. The electrode contacts were arranged in a staggered grid, with a pitch of 45 µm in both lateral dimensions and a nearest-neighbor center-to-center distance of 31.8 µm.

A final 250-nm PI layer was then applied over the device to protect the IrOx contacts during subsequent processing. Openings in the PI above the bonding pads and electrode contacts were defined by oxygen-plasma etching using a PR1-7000A1 (Futurrex, Inc.) photoresist mask. The photoresist mask and processing residues were subsequently removed using acetone, Remover PG and Remover RR-41. An additional 5-nm Ti/100-nm Au layer was then deposited over the BGA pads using the same BGA-pad mask to improve solder wetting during device bonding. The resulting probes had an overall thickness of approximately 2–5 µm, depending on device design.

The wafer was then diced, solder balls were placed onto the BGA pads, and individual devices were bonded to printed flexible cables. The bonded BGA interface was then underfilled with epoxy. The probe was subsequently released from the silicon substrate by wet-etching the underlying Ni sacrificial layer and connected to a data-acquisition system (SpikeGadgets or Intan) for impedance measurements. The loop at the distal end of the probe was manually threaded onto the implantation shuttle and the probe was fixed in place using polyethylene glycol with molecular weight of 1K. Assembled devices were sterilized using room-temperature ethylene oxide process (Anprolene AN74i).

### Wafer-level impedance and electrochemical characterization

The electrode impedance was measured at 1 kHz in phosphate-buffered saline (PBS) to evaluate channel yield. The median impedance of a typical device was 80 – 200 kΩ. Functional channels were defined for each device as those with impedance 0.7 – 1.5 times its median. Device-specific functional-channel yield was the number of channels in this range divided by the total channel number as in Figure S4A, which presents the impedance distribution for 100 devices from 3 representative wafers. Median selected-device yield was 91.4% (interquartile range, 86.4–95.3%). Box limits indicate the first and third quartiles, center lines indicate medians and whiskers extend to the most extreme observations within 1.5 times the interquartile range. Individual lower and upper out-of-range values are shown as boundary triangles.

Electrochemical impedance spectroscopy (EIS) was performed on one representative IrOx electrode before and after cumulative biphasic pulsing at 10 μA (Figure S4B). The cathodic-first, charge-balanced cycle comprised 33.4 μs at zero current, -10 μA for 100 μs, 66.7 μs at zero current, +10 μA for 100 μs and 1.5 ms at zero current, giving a cycle duration of 1.8 ms (555.52 cycles s⁻¹). EIS was repeated after 1, 2, 10, 35, 45 and 65 million cumulative pulses. Potentiostatic spectra were acquired from 1 Hz to 1 MHz at 10 points per decade using a 10-mV rms perturbation. The measured point nearest 1 kHz was identified in each spectrum. These were sequential measurements of one electrode and were therefore summarized descriptively without inferential statistics.

### Analytical estimates for EBL interconnect design

For Figure S1A, gold-interconnect resistance was calculated as R = ρ(w)L/(wt), where L is trace length, w is linewidth, and t = 100 nm is the gold thickness. Linewidths of 50-1,000 nm and trace lengths of 5, 10, 15 and 20 mm were evaluated. The linewidth-dependent resistivity model used a bulk resistivity ρ0 of 2.2 × 10⁻⁸ Ω m, an electron mean free path of 37.7 nm, a 12.9-nm dimensional offset and a scale factor of 1.09. Defining k = min[(w -12.9 nm)/(2 × 37.7 nm), 1], resistivity was calculated as ρ(w) = 1.09ρ0/[k(1 -ln k)]^42^.

For Figure S2A, ideal electron-beam exposure time for 100 parallel, 10-mm-long interconnects was calculated as dose × line length × linewidth × interconnect number divided by beam current, using a dose of 250 μC cm⁻². Beam currents were 100 nA for linewidths >500 nm, 50 nA for linewidths >250-500 nm, 10 nA for linewidths >100-250 nm and 4 nA for linewidths of 50-100 nm.

Mean time to failure (MTTF; Figure S2B) was estimated using an entropy-production model^44^. Calculations used a temperature of 300 K, a 10-mm trace length, a 100-nm gold thickness, a gold atomic concentration of 5.9 × 10²⁸ m⁻³, an effective charge number of 11, a resistivity of 2.6 × 10⁻⁸ Ω m, an atomic volume of 1.25 × 10⁻²⁹ m³, a critical void dimension of 0.2 nm and a vacancy formation energy of 1.6 × 10⁻¹⁹ J. Gold diffusivity at 300 K was estimated as 5.23 × 10⁻²⁴ m² s⁻¹ by scaling a value of 10⁻²⁰ m² s⁻¹ at 573 K with an activation energy of 0.41 eV. Current density is the applied current divided by the cross-sectional area (100 nm × linewidth).

### Crosstalk simulation

Electrical crosstalk between neighboring interconnects was evaluated using a combined finite-element and circuit-simulation workflow. First, two-dimensional electrostatic / electric-current finite-element models were constructed in COMSOL to extract per-unit-length coupling and shunt parameters for adjacent gold interconnects embedded in polyimide and surrounded by phosphate-buffered saline (PBS). Both in-plane and out-of-plane neighboring trace geometries were considered. The COMSOL models used an enlarged PBS domain to approximate an infinite bath boundary, with refined meshing around the metal traces and polyimide/PBS interface. For each geometry, the simulations extracted mutual coupling capacitance between traces, shunt capacitance from each trace to the surrounding medium, and conductance terms representing leakage through the dielectric/electrolyte environment.

The extracted COMSOL parameters were then converted into a distributed two-line LTspice circuit model. Each trace was represented as a ladder of 24 sections over a total probe length of 10 mm. Each section included series interconnect resistance, shunt capacitance and conductance to ground, and mutual capacitance and conductance between the aggressor and victim traces. The line resistance was calculated from trace geometry using a 100 nm gold thickness and bulk gold resistivity. The polyimide thickness used for the parameter-sweep simulations were 2 µm.

For each LTspice simulation, a 1 V AC input was applied to the aggressor line through the electrode-site impedance (Z_site_), while the victim line was driven with 0 V AC through the same impedance. Both far ends were terminated with a 13 MΩ load. The simulations used a conservative Z_site_= 2 MΩ, and polyimide conductivity of σPI = 1 × 10^−9^ S/m was assumed as a conservative estimate to account for potential dielectric leakage under chronic exposure to saline, and were evaluated at 1 kHz. Crosstalk was defined as 100 × |V_victim,far_ / V_in_ |, and signal transfer was defined as |V_aggressor,far_ / V_in_ |. Signal attenuation was calculated as 100 × (1 -|V_aggressor,far_ / V_in_ |).

The parameter sweeps varied interconnect linewidth, gap-to-linewidth ratio, and neighboring-trace orientation. For the large-sweep panels, linewidths ranged from 1 nm to 100 µm and gap-to-linewidth ratios were 0.2, 0.5, 1, 2, and 4. For the design-relevant condition of 180 nm linewidth and 220nm gap, interpolation between the simulated points gave a nominal crosstalk of approximately 0.8% and signal attenuation of approximately 13.4% for a 10 mm probe.

### Animal Surgery

The chronic implantation procedure followed methods described in previous studies^12^. Briefly, animals were kept under anesthesia throughout the procedure, starting at 3–5% and maintained at 1–2% isoflurane, depending on body weight. A 2–3 mm diameter craniotomy window for mice, or a smaller window for rat, was created for implantation. For mice, the dura was left intact, eliminating the need for removal, while for rats, a small opening was made in the dura to facilitate implantation, minimizing surgical damage to brain tissue.

The probe was guided using a tungsten shuttle with a sharpened T-shaped tip at an implantation speed of 10–100 µm/s following an initial puncture through the dura. Upon reaching the target depth, PBS was applied to dissolve the polyethylene glycol (PEG) and provide moisture to the brain surface, a process that took approximately 5 minutes.

After retracting the shuttle, the probe was left in place, and a stereotactic device was used to gently position it, leaving slight redundancy above the brain surface. A glass window was placed to seal the area for the 2-3mm cranial openings, ensuring that cerebrospinal fluid (CSF) or saline filled the volume to maintain secure contact between the neural probe and the skull.

The glass window was secured, or the small cranial opening was covered, using Kwik-Sil (World Precision Instruments) and super glue (Loctite). A stainless-steel wire was implanted as a grounding/reference wire. The probe assembly was fixed to the skull with dental cement (C&B-Metabond), and a stainless-steel head bar was attached above the cement to facilitate head fixation during awake electrophysiology recordings. For rat surgeries, a screw was implanted into the skull to contact the CSF. The electronics setup, including a SpikeGadgets FPGA, amplifier, routing board, and terminal board, was stacked and encapsulated within a customized 3D-printed cage. The electronics were grounded to the screw using a metal wire.

### Electrical stimulation and recording

Each EBL-NET contact was capable of both recording and electrical stimulation through its IrOx-coated electrode surface. When interfaced with compatible electronics, the probes supported concurrent stimulation and recording. Selected contacts were stimulated individually in randomized sequential trials using a Ripple stimulation system. Charge-balanced biphasic pulses were delivered at amplitudes of 1–5 μA. The cathodic and anodic phases were each 167 μs in duration and were separated by a 67-μs interphase interval. Each stimulation train consisted of 50 pulses delivered at either 50 or 100 Hz, depending on the session, corresponding to intervals of approximately 1 or 0.5 s, respectively, between the first and last pulses. Each train was followed by a 2.5-s stimulation-free interval, and each combination of stimulation contact and current amplitude was repeated ten times per session. To reduce amplifier saturation and facilitate recovery after stimulation, a 1-ms channel reset was applied following each pulse. Stimulation contacts were selected from a subset of contacts associated with sorted unit activity during recording.

Electrical stimulation produces transient recording artifacts arising from electrode polarization, stimulus coupling and amplifier recovery. To reduce their effects on spike detection and sorting, data from 0.4 ms before to 2.4 ms after each stimulation pulse were excluded from analysis. This blanking procedure necessarily precluded analysis of spikes occurring within the excluded interval, including short-latency responses that could reflect direct neuronal activation. It nevertheless improved spike-sorting robustness and enabled analysis of neuronal activity occurring outside the artifact window, including sustained and longer-latency stimulus-evoked responses.

### Chronic neural recording and behavior studies in rats

The first rat was implanted in the anterior cingulate cortex (ACC) at coordinates 3.0 mm AP, -0.6 mm ML, and 2.5 mm DV relative to bregma. This rat was trained to perform a binary action-sequence task in a box with high plastic walls and a stainless-steel floor measuring 25 cm by 34 cm^64^. Nose ports were arranged along one 25 cm wall. A custom-programmed microcontroller controlled and monitored the lights, nose ports, and reward delivery system, and interfaced through USB with a PC running a MATLAB-based control program. The central initiation port contained a white LED that signaled the start of a new trial (action onset). After trial initiation, the central LED was extinguished, and LEDs above the left and right side ports were illuminated to indicate that the side ports were available for selection. Side ports delivered liquid reward (0.1 ml drops of 10% sucrose mixed with black cherry Kool-Aid) through a motorized syringe pump when the rat selected the ports in the trained sequence, which was left-left-right (L-L-R) in this experiment. Neural recordings began 115 days after implantation and continued until day 180. Spike sorting was performed using Kilosort4.

The second rat was implanted to record neural activity from hippocampal CA1 and dentate gyrus (DG) at coordinates -4.0 mm AP, -2.0 mm ML, and 4.5 mm DV relative to bregma. Before surgery, the rat was trained to shuttle between two food wells positioned at opposite ends of a linear track. To receive food reward, the rat was required to traverse the full length of the track rather than obtain food from only one end. Training continued until the rat reached a standardized performance criterion.

Neural recordings began 90 days after implantation and continued across multiple sessions until day 104. During each session, a camera recorded the rat’s trajectory along the track, allowing position to be correlated with neural activity to identify place cells in CA1. Spike sorting was performed using MountainSort4.

### Intraoperative human recording

Two EBL-NET probes were acutely implanted into the left anterior middle temporal gyrus in tissue scheduled for resection during a left anterior temporal lobe resection for medically refractory epilepsy. Podcasts were played while the patient was under general anesthesia. Each flexible probe was delivered with a laser-machined tungsten shuttle wire, and extracellular recordings were acquired before and after shuttle-wire retraction. Session 1 was recorded before shuttle-wire retraction, and Session 2 was recorded after retraction. Recordings were acquired at 30 kHz with Open Ephys from two 128-channel probes. Channels identified as bad were excluded before analysis.

### Histology and immunohistochemistry

Following completion of the chronic implantation experiment, animals were deeply anesthetized with isoflurane (3%-5%) and transcardially perfused with phosphate-buffered saline (PBS) followed by 4% paraformaldehyde (PFA); both fluids were chilled to 4 °C. Following perfusion, the head was removed from the body and stored in 4% PFA for 48 hours. Care was taken to remove the head from the brain and maintain the probes intact. The brains were then cryoprotected in a 10% sucrose solution for 24 hours. Tissue containing the implanted 128-channel EBL-NET probes was sectioned perpendicular to the probe trajectory into 30-µm-thick sections using a cryostat (CM1520 Leica Biosystems, IL). Histological preparation was adapted from previously reported protocols for NETs.

Tissue slices were prepared for histology by first rinsing the tissue three times in a 1 × PBS solution with 0.1% triton X-100 for 5 minutes each before blocking the tissue in a 10% BSA solution for 1 hour with gentle agitation at room temperature. Slices were then incubated with primary antibodies overnight at 4°C for astrocytes (mouse anti-GFAP, 1:250, Millipore MAB3402), microglia (rabbit anti-Iba1 1:200, Fujifilm 019–19741), and neurons (NeuroTrace 640/660 1:200, Invitrogen). The slices were then washed with PBS and incubated with secondary antibodies for 2 h at room temperature (donkey anti-mouse 488, 1:100, Invitrogen A-21202; donkey anti-rabbit 594, 1:100, Invitrogen A-21207). After staining, the slices were mounted on a glass coverslip with Vectashield with DAPI before imaging. Bright-field and fluorescence images were acquired from the corresponding sections to identify the probe location and tissue morphology.

## Quantification and statistical analysis

### Spike sorting, longitudinal neural tracking, and chronic stability quantification in mice

Spike sorting was performed to isolate single units from the raw 30kHz electrophysiological data collected by the EBL-NETs. We employed the SpikeInterface^46^ and MountainSort 5 algorithms (scheme 2, threshold 5.5) unless otherwise mentioned for time interval segmentation, feature extraction, clustering, and post-processing, followed by manual curation based on unit waveform distribution across neighboring channels, firing rates, auto-correlograms (ACGs) and cross-correlograms (CCGs) for noise rejection, unit de-duplication, and oversplit units merging.

For 10-day longitudinal neural tracking for mouse 1 (Table S1, Figures 2D–2H), electrophysiology recordings were conducted for 30 minutes each day at the same time of the day for 10 consecutive days. The recordings were concatenated and processed for unit sorting, then the units assigned back to each day. Using this method, the same neuron was assigned a consistent identifier across all 10 days.

For unit tracking quality control, thresholds were applied for signal-to-noise ratio (SNR), firing rate, waveform similarity, and average unit location variations, calculated as the center of mass of the amplitude distribution. Units that did not meet these criteria on more than half of the days were excluded from further analysis. The units passed the quality check were used to evaluate the stability of recordings over the 10-day period.

For recording stability quantification, we summarized 12 recording sessions spanning days 13-176 after implantation (Figure S6). For each unit-session waveform set, the median per-unit spike maximum-channel peak-to-peak amplitude was calculated.

Individual unit values, the session median and the interquartile range are shown. Electrode impedance was measured at 1 kHz for 128 channels per session and is displayed as individual channel values with the session median and interquartile range on a logarithmic axis.

### Putative monosynaptic connection detection

Putative monosynaptic connections were identified following previously described methods^61–63^. For baseline recording data, we computed the cross-correlogram (CCG) for each neuron pair, focusing on bins near 0 with a bin size of 0.5 ms and an analysis window spanning -4 ms to 4 ms to detect peaks significantly different from baseline. To assess significance, spike timestamps for both units were jittered by randomly shifting them within a window of (-10 ms, 10 ms), and this process was repeated 1,000 times to generate a distribution of jittered CCGs. A connection was considered putative if both units exhibited firing rates greater than 1 Hz, the peak within ±4 ms exceeded the 99% confidence interval (CI) of the jittered distribution, and the peak value had a Z-value greater than 3 when compared to the baseline distribution.

### Stimulation response and connectivity analysis

Neural responses were assessed separately for each combination of recorded unit, stimulation contact and stimulation current. For activation mapping and cross-animal analyses, firing rates during each of ten stimulation trains were compared with firing rates during the corresponding 2-s pre-train baseline intervals. A response was classified as activated when the mean firing rate during stimulation exceeded the mean baseline firing rate and a two-sided Welch’s t-test yielded (P ≤ 0.005). For the representative current-response analyses in Figures S8A and S8B, mean and s.e.m. were calculated across ten paired trials at each current. For the cross-animal analysis in Figure S8C, measurements from all 10 stimulation contacts in each of two sessions were retained at every tested current for each of five animals. Thus, the mean and s.e.m. for each animal and current were calculated from 20 contact-session observations.

Population activity patterns associated with stimulation through different contacts were quantified using stimulation-period firing rates across simultaneously recorded units. For each trial, the resulting neuron-dimensional firing-rate vector represented the population activity state. Uniform Manifold Approximation and Projection (UMAP) was used to project these high-dimensional firing-rate vectors into three dimensions for visualization (Figure 3F).

For comparisons of stimulation-associated connectivity patterns in Figure S9, ten neighboring pairs of stimulation contacts were used to compare connectivity-source units and directed source-target edges identified during stimulation through contacts A and B. Entities shared between the two stimulation conditions were defined by the set intersection, whereas contact-specific entities were defined by the corresponding directional set differences. The union comprised all entities observed under either stimulation condition.

For chronic stimulation experiments, data from all sessions were concatenated after stimulation-artifact removal and subjected to spike sorting. The resulting units were subsequently separated according to their individual recording sessions for longitudinal analysis. As in the longitudinal unit-tracking analysis, unit quality and stability were evaluated across days using signal-to-noise ratio (SNR), firing rate, waveform similarity and changes in estimated unit location. Unit location was estimated from the center of mass of the spike-amplitude distribution across recording contacts. Units that failed the predefined quality or stability criteria on more than half of the recording days were excluded from subsequent analyses; the remaining units were retained for longitudinal evaluation.

In Figure S10A, we plotted the spike waveforms across neighboring channels for a representative unit tracked across the seven sessions. For the pulse-aligned analysis in Figure S10B, spikes from the unit were aligned to 500 stimulation pulses delivered through the same contact in each of seven recording sessions. Spikes occurring from +2.4 to +9.6 ms relative to pulse onset were binned at 0.25-ms resolution, and bin counts pooled across pulses were smoothed using a Gaussian kernel with a SD of 0.5 ms. The interval from −0.4 to +2.4 ms relative to stimulation onset was designated as the stimulation-artifact interval. The plotted trace represents the smoothed pooled spike count.

For the longitudinal stimulation analyses in Figures S11–S12, the activation threshold was defined as the lowest tested stimulation current satisfying the predefined activation criterion. Session-level measurements of activation threshold and firing-rate (FR) modulation (defined as (FR during stim – baseline FR) / (FR during stim + baseline FR, where baseline is the 2s recording before stimulation onset) were grouped according to recorded unit and stimulation contact, yielding 51 unit–stimulation-contact pairs. In panel a, bars and error bars represent the arithmetic mean and sample standard deviation, respectively, across the three to seven sessions in which each pair was detected; individual points represent session-level observations. In panel b,c, we report the mean and SD distributions for quantities (activation threshold and firing rate modulation) of the stimulation channel -unit pairs.

### Rat unit tracking, population decoding, and place-cell analysis

ACC activity was analyzed across four recording sessions corresponding to post-implantation days 115, 116, 120, and 180. For the example sequential population activity heatmap, spike trains were binned at 10 ms and convolved with a causal one-sided Gaussian kernel (sigma = 100 ms, truncated at 5 sigma and normalized to unit sum) to estimate instantaneous firing rates. Activity was aligned to action onset from -2 to +2 s. Units with mean firing rate below 0.25 Hz in the displayed action-aligned window were excluded. Firing-rate traces were z-scored for each unit across the concatenated action-aligned matrix, and units were reordered using RasterMap^48^ to group units with similar temporal activity patterns.

For the representative single-unit firing-rate example, mean firing-rate traces were computed separately for left and right actions across the four recording sessions. Shaded regions indicate the s.e.m. across trials. Each day’s recording session was sorted, and stable ACC units were identified based on the consistency of their waveforms, amplitudes and ISI distribution across sessions. For the single-unit decoding stability analysis, left-versus-right decoding scores were computed for stable units across the four sessions, and adjacent-session Pearson correlations were calculated across unit decoding scores.

For the ACC population decoding analysis, action-level firing vectors were constructed from matched stable units for each step in the L-L-R sequence. Each element of a population vector was calculated by the firing rate of a single unit from 0-500ms after an action onset. Each action was represented as one population vector in neural state space. Principal component analysis (PCA) was fit to the day-115 action-by-unit firing matrix without per-neuron firing-rate normalization. The resulting day-115 PC coefficients were then used to project all later sessions into the same two-dimensional neural state space. A three-class linear discriminant analysis (LDA) classifier was fit using day-115 PC1 and PC2 scores to classify the first L, second L, and R sequence steps. The day-115 decision boundaries were applied to later sessions to evaluate stability of the population geometry. Boundary-shifted decoding was also evaluated after translating the day-115 LDA boundaries by the session-wise population mean shift in PC space.

For the representative hippocampal place-cell example, the same unit was tracked across sessions on post-implantation days 90, 91, 92, 97, and 104 by sorting the recording concatenated from those recorded in those five days. Mean waveforms, interspike-interval distributions, and occupancy-normalized spatial firing-rate maps were plotted for each session. Unvisited or masked spatial bins were rendered as white pixels in the spatial firing-rate maps.

### Intraoperative drift and unit analysis

For drift estimation and visualization, voltage traces were band-pass filtered from 300 to 6000 Hz, and globally median common-referenced. Locally exclusive negative threshold-crossing events were detected with SpikeInterface 0.102.3 using a threshold of 7, a 0.8-ms exclusion window, and an 80-µm detection radius. Detected events were localized by monopolar triangulation. Nonrigid motion was then estimated with DREDge^54^ from the localized event-density image by computing pairwise depth shifts over time and solving for displacement in four Gaussian windows spaced by 400 µm along the probe depth axis.

The drift panel in Figures 5D and 5E shows EBL-NET-2 before and after shuttle-wire retraction. This window contained 2,591 localized threshold-crossing events before retraction and 5,374 events after retraction. Events were plotted at their detection times and localized depths, with vertical ticks colored by absolute peak amplitude on a shared grayscale scale. Local DREDge displacement traces were plotted in gray and their mean in blue.

The representative single-unit map uses the post-retraction EBL-NET-2 recording and shows the 100 representative semantic-analysis units used for the corresponding semantic-encoding analysis denominator. Units labeled as noise were excluded, and merge groups were represented by their retained representative unit. Unit locations were taken from the curated localization results. Bubble area was scaled by the peak-to-peak waveform amplitude on each unit’s largest-amplitude channel, and representative local waveform groups were plotted with electrode contacts and unit locations overlaid.

### Lexical-semantic encoding analysis

Semantic encoding was analyzed using a passive-speech listening workflow adapted from prior human single-neuron language studies^55,56^. For each presented word, Word2Vec embedding vectors were reduced by principal component analysis, and the first 10 principal components were retained. Spike counts were summed over each word duration for each neuron.

For each unit, word-evoked spike counts were modeled with a ridge-regularized Poisson generalized linear model with a log link. Predictors included the 10 Word2Vec principal-component scores, word duration, and the interaction between each principal component and word duration, yielding 21 z-scored predictors per word. Model performance was evaluated with a five-iteration nested training procedure. In each iteration, data were split into 80% training and 20% held-out test sets. Within the training set, fivefold cross-validation selected the regularization strength by optimizing cross-validated log likelihood, and the selected model was evaluated on the held-out test set.

Statistical significance was assessed with matched null models generated by shuffling embeddings across words 100 times and refitting the model. A neuron was considered semantically encoded when the observed model significantly exceeded the shuffled-model distribution by permutation testing, the observed-minus-shuffled median log-likelihood difference was positive, and McFadden’s pseudo-R2 was positive.

For the figure-level summary, each neuron was counted as significant or not significant for semantic encoding. The proportion of significant neurons was compared with a 5% false-positive null using an exact one-sided binomial test. Error bars show exact two-sided 95% Clopper-Pearson confidence intervals. Before-versus-after aggregate comparisons were performed with two-sided Fisher exact tests. Significant semantic encoding exceeded the 5% null in both probes before and after shuttle-wire retraction.

### Declaration of generative AI and AI-assisted technologies in the manuscript preparation process

During the preparation of this work, the author(s) used OpenAI to assist with grammar checking and language polishing. The author(s) reviewed and edited the output as needed and take full responsibility for the content of the published article.

