## supplementary information for "Nanofabricated ultraflexible electrodes for chronic, high-density neural recording and stimulation"

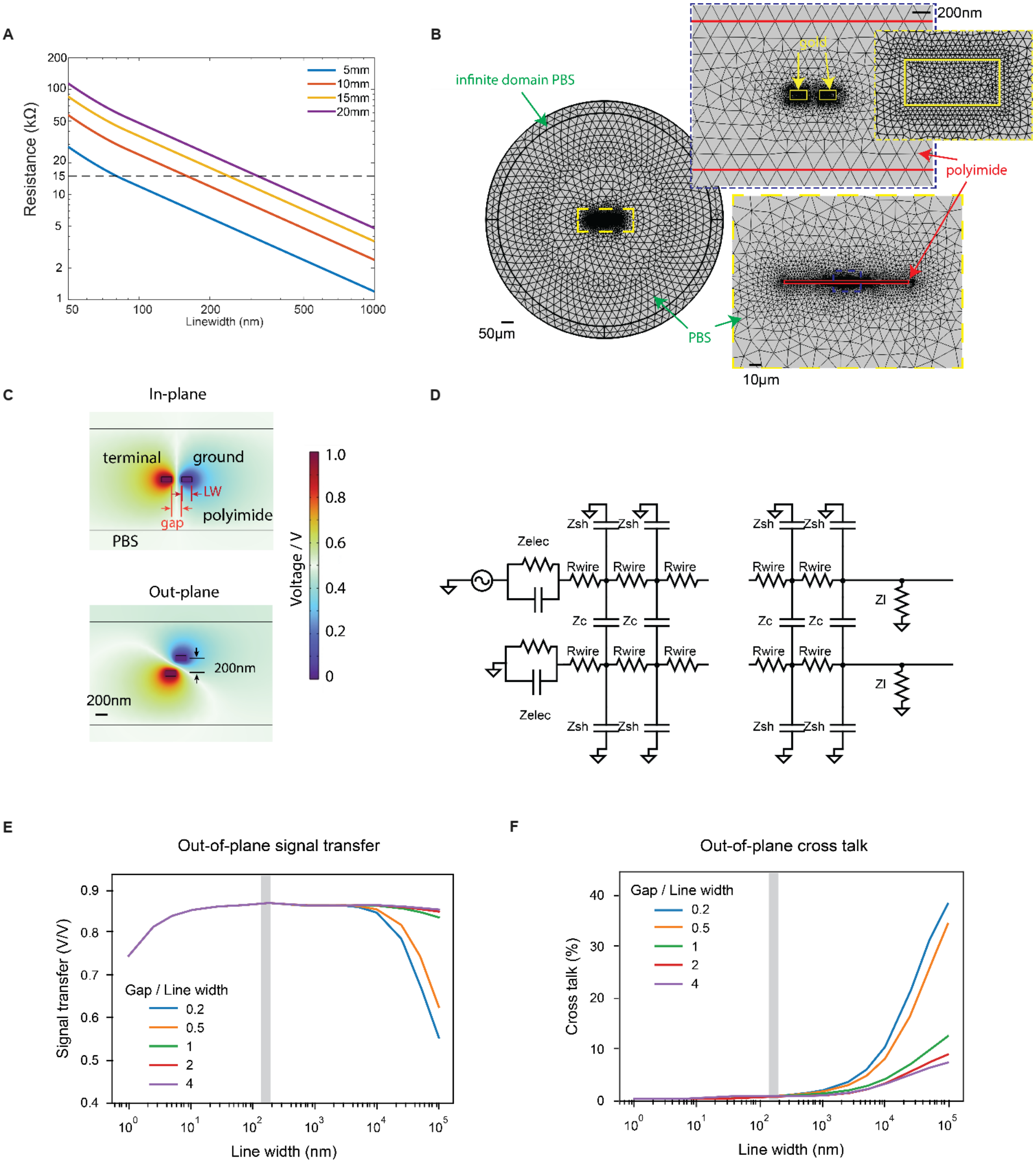


**Figure S1. Electrical modeling for interconnect resistance, crosstalk, and signal transfer in densely routed EBL-NETs.** (A) Calculated resistance of 100-nm-thick gold interconnects as a function of linewidth for trace lengths of 5 – 20 mm; the dashed line marks the 15 kΩ design threshold. (B) COMSOL finite-element model to extract coupling and shunt parameters, showing the infinite-domain PBS boundary, local mesh, and polyimide/PBS geometry around neighboring traces. (C) Representative COMSOL voltage distributions for in-plane and out-of-plane neighboring interconnect geometries. (D) Distributed two-line LTspice circuit used to simulate aggressor line and victim lines, including interconnect resistance (Rwire), mutual coupling (Zc), shunt paths (Zsh), electrode-electrolyte interface impedance (Zelec), and amplifier/input impedance (ZI). (E) Out-of-plane signal transfer (Vout/Vin) as a function of line width for gap-to-linewidth ratios of 0.2, 0.5, 1, 2, and 4. (F) Out-of-plane crosstalk as a function of line width for the same gap-to-linewidth ratios. For (E) and (F) σPI = 1 × $10^{-9}$ S/m, trace length = 10 mm, metal thickness =100 nm, PI thickness = 2 μm, contact site impedance = 2 MΩ, and 24 LTspice ladder segments.

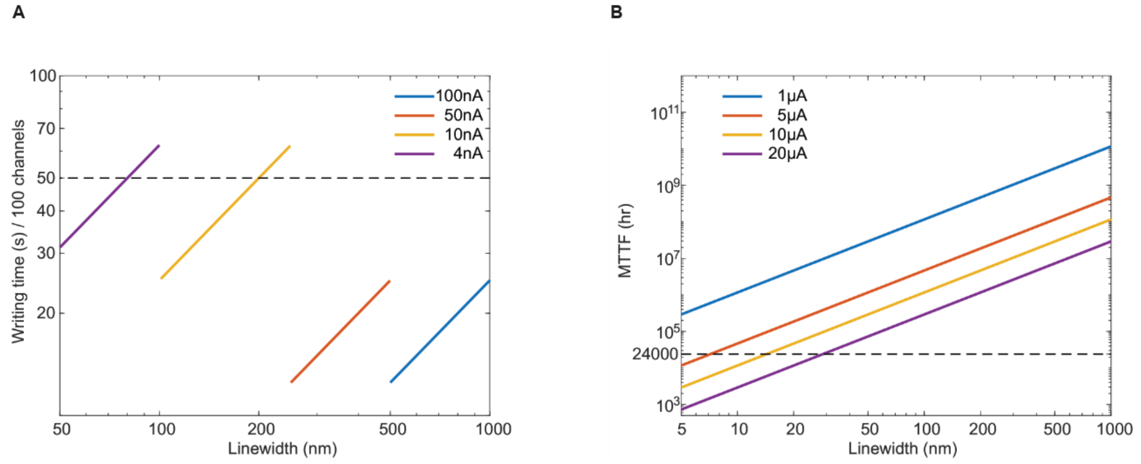


**Figure S2. EBL fabrication and reliability considerations for nanoscale interconnects.** (A) Estimated electron-beam exposure time required to write 100 parallel 10-mm-long interconnects as a function of linewidth and beam current, assuming an exposure dose of 250 μC $cm^{-2}$. The dashed line marks 50 s. (B) Estimated mean time to failure (MTTF) of 10-mm-long, 100-nm-thick gold interconnects as a function of linewidth at applied currents of 1, 5, 10, and 20 μA. The dashed line marks 24,000 h, approximately 1,000 days.

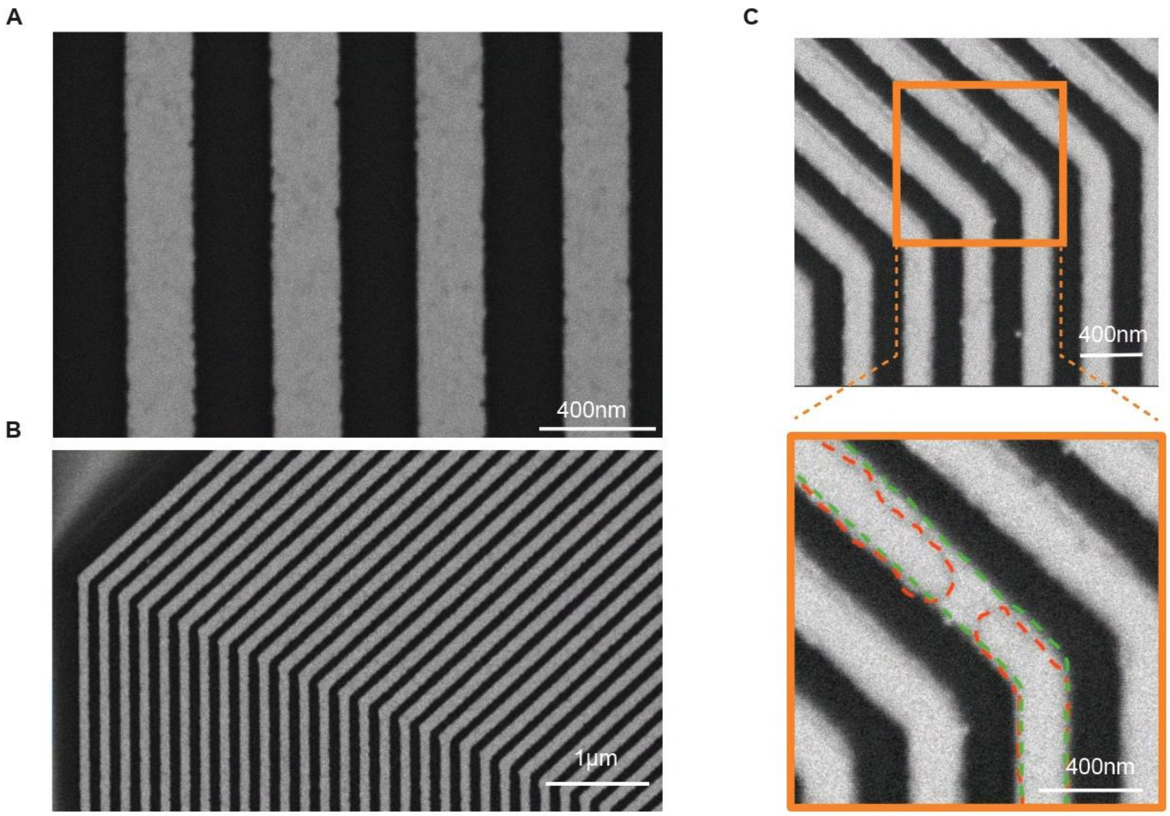


**Figure S3. SEM characterization of the EBL-patterned interconnects.** (A) High-magnification SEM image of 200nm-wide EBL-patterned lines. (B) Angled interconnect turns showing clean lift-off and precise double-pass alignment along both axes. (C) Example of a trace disrupted after the first metal pass and restored by double-pass writing. Inset: red and green outlines indicate the first- and second-pass metal patterns, respectively. The second-pass metal bridges the defect and restores electrical continuity.

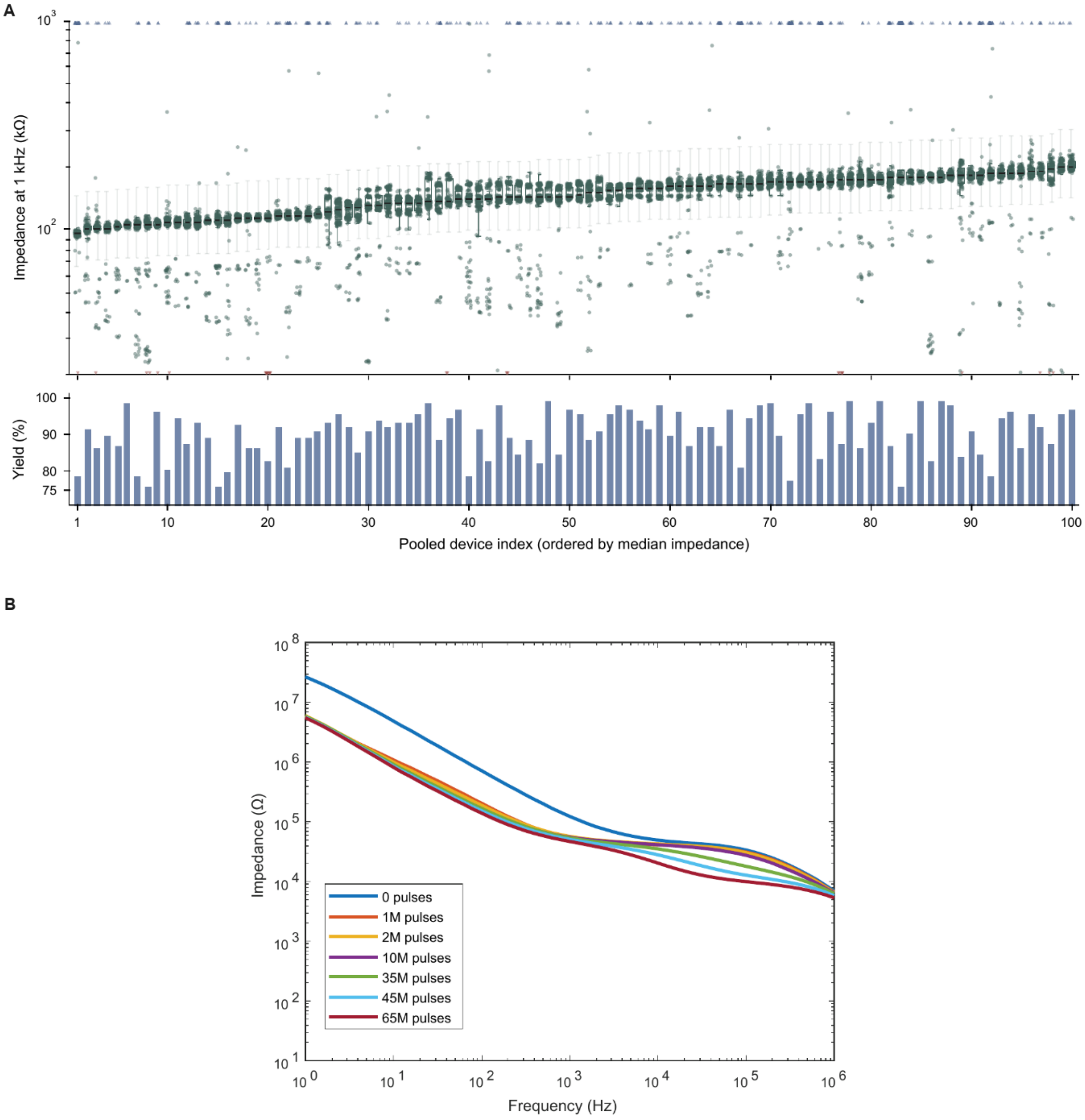


**Figure S4. Device-level yield and electrochemical impedance stability.** (A) Device-level 1-kHz impedance distributions (top) and corresponding functional channel yields (bottom) for devices pooled across three wafers. Devices median impedances ranged 80 – 200 kΩ, with functional channels defined as those within 0.7–1.5× the device median. Boxes indicate the first and third quartiles (Q1–Q3) with median and 1.5× interquartile-range (IQR) whiskers; points indicate individual channels. Triangles denote measurements outside the displayed 20–1,000 kΩ range. (B) Electrochemical impedance spectrum of a representative IrOx contact measured before stimulation and after 1, 2, 10, 35, 45, and 65 million cumulative biphasic pulses at 10 μA.

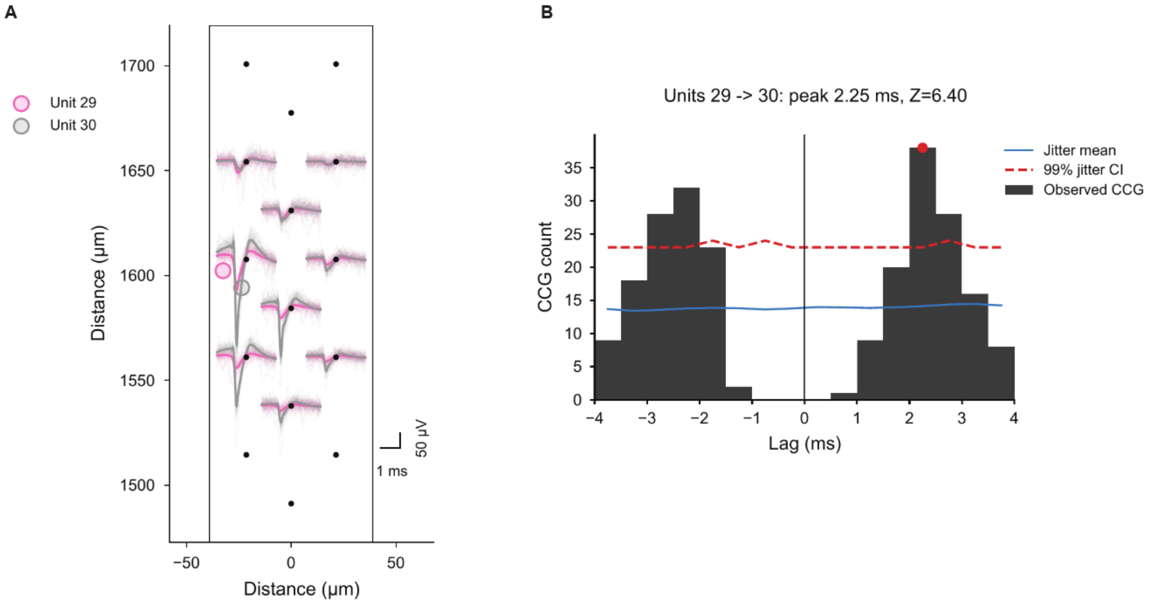


**Figure S5. Waveform and spike-train validation of a putative short-latency functional connection.** (A) Spatial waveform distributions of units 29 and 30 on neighboring contacts. Faint traces show 40 deterministically sampled waveforms per unit, bold traces show mean templates, black points mark recording contacts, and colored circles mark estimated unit locations. (B) Cross-correlogram (CCG) for units 29 and 30. Bars show the observed CCG, the blue line shows the mean of the jittered CCG distribution, and the red dashed line shows its pointwise 99th percentile. The largest peak occurred at +2.25 ms, indicating a putative connection from 29 to 30. Additional significant bins near -2.25 ms indicate a putative connection in the reversed direction, consistent with bidirectional functional coupling between the pair.

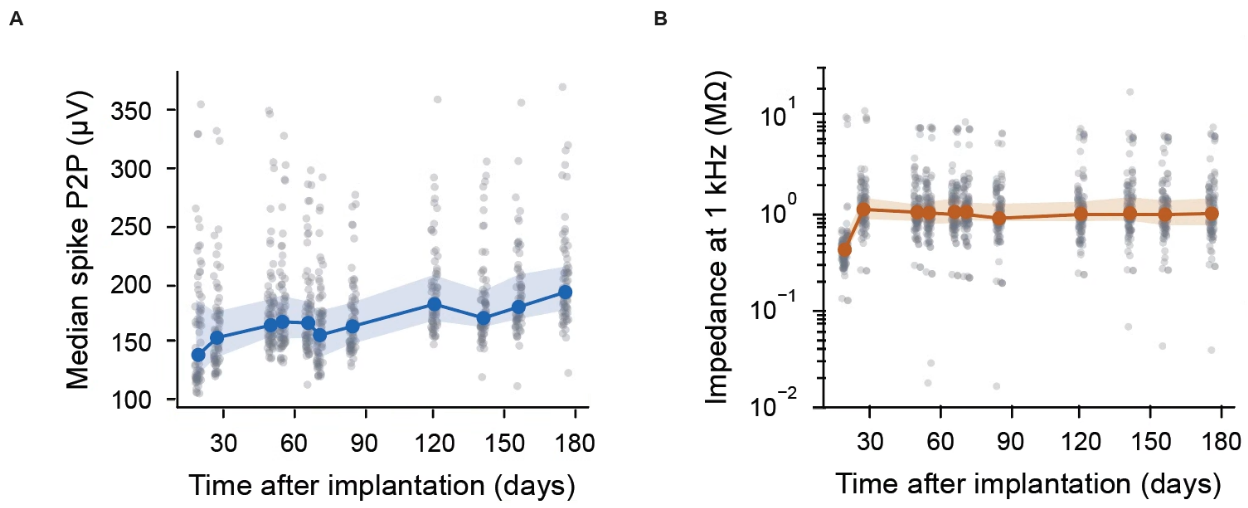


**Figure S6. Six-month recording stability of the implanted EBL-NETs**. (A) Median spike peak-to-peak (P2P) amplitude across longitudinal recording sessions. Gray points show individual unit-session values, the line shows the session median, and shading shows the interquartile range; recording time is plotted as elapsed days after implantation. (B) Electrode impedance at 1 kHz across the same longitudinal period. Gray points show individual channels, the line shows the session median, and shading shows the interquartile range; impedance is plotted on a logarithmic axis.

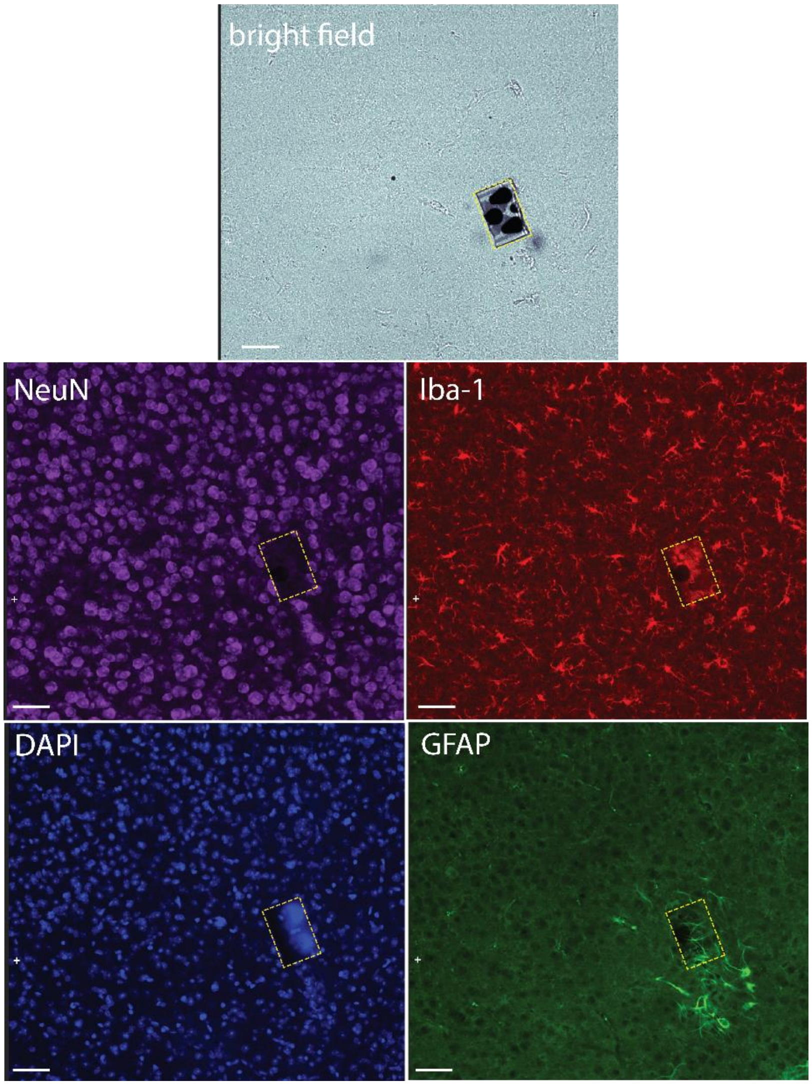


**Figure S7. Histological characterization of brain tissue surrounding a chronically implanted EBL-NET 4 months after implantation.** Representative bright-field and fluorescence micrographs of a 30-μm-thick brain section containing the implanted EBL-NET. Immunofluorescence staining labels neurons (NeuN, magenta), microglia (Iba-1, red), cell nuclei (DAPI, blue), and astrocytes (GFAP, green). The EBL-NET is indicated by the dashed box. Scale bars, 50 μm.

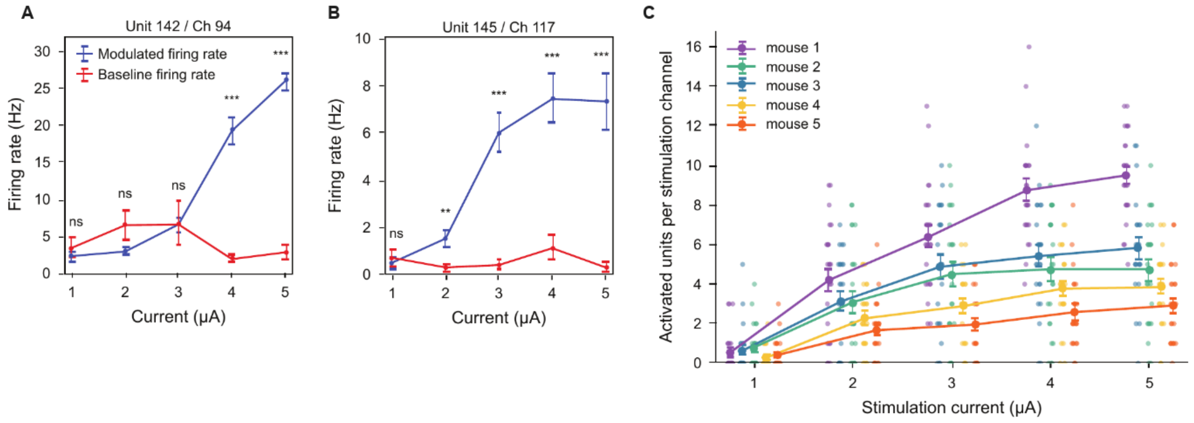


**Figure S8. Current-dependent activation by EBL-NET ICMS.** (A, B) Representative current-dependent spiking responses of isolated units evoked by 1–5 μA pulse trains delivered through a single contact. Blue: firing rates during stimulation; red: duration-matched pre-stimulation baseline. Dots show the mean of ten trials and error bars indicate s.e.m. Significant activation was detected at 4 and 5 μA in (A) ($p = 0.379$, 0.108, 0.975, $8.94 \times 10^{-6}$, and $2.85 \times 10^{-9}$ at 1–5 μA, respectively), and at 3–5 μA in (B) (Two-sided paired t-tests; $p = 0.642$, 0.00863, $1.26 \times 10^{-4}$, $7.99 \times 10^{-4}$, and $1.36 \times 10^{-4}$ at 1–5 μA, respectively). (C) Number of activated neurons as a function of stimulation current from 1 to 5 μA (n=5 mice). Small colored points represent individual stimulation-channel/session response. Matching colored lines and error bars show the animal-specific mean ± s.e.m. across 20 responses per animal and current (10 channels × 2 sessions).

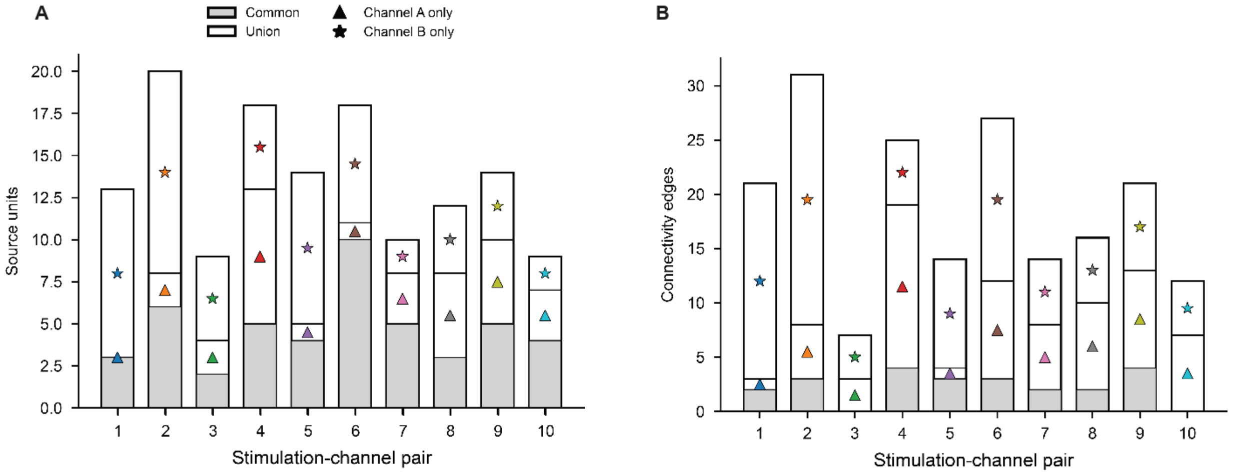


**Figure S9. Shared and stimulation-site-specific functional coupling between nearby stimulation pairs.** Stimulation was delivered through 10 neighboring channel pairs to compare connectivity-associated source units and putative connectivity edges shared by, or specific to, each stimulation site. (A) Source units participating in putative connections for each pair. Gray bar shows units common to both stimulation conditions. Outlined bar shows the union, with an internal divider separating channel-A-only and channel-B-only units; colored triangles and stars identify the respective channel-specific segments. (B) Putative monosynaptic connectivity edges for the same pairs, using the same common, channel-A-only, channel-B-only, and union encoding.

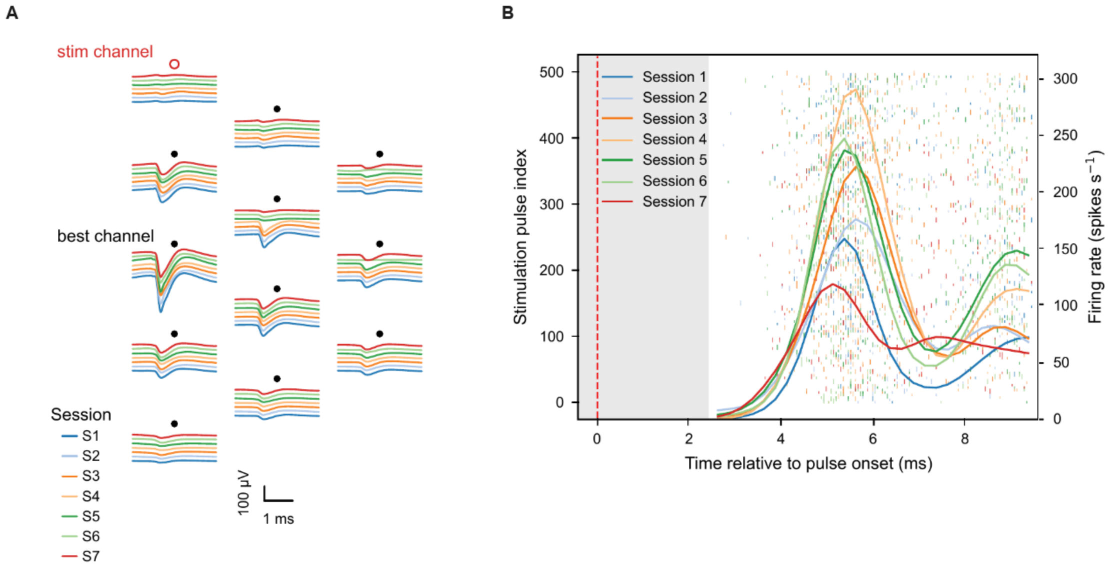


**Figure S10. Chronic stability of stimulation-evoked unit responses.** (A) Waveforms of a representative unit across seven ICMS sessions spanning two weeks. Waveforms from each session are color-coded and vertically offset to show the stability of the spatial waveform footprint. The stimulation contact is marked by a red open circle, while neighboring recording contacts are indicated by black dots. (B) Spike rasters (vertical bars) and Gaussian-smoothed pooled firing rates (curve) from the same unit aligned to 500 stimulation pulses per session. Red dashed line marks stimulation onset and the gray window marks the -0.4 to 2.4 ms blank-out interval due to stimulation artifacts.

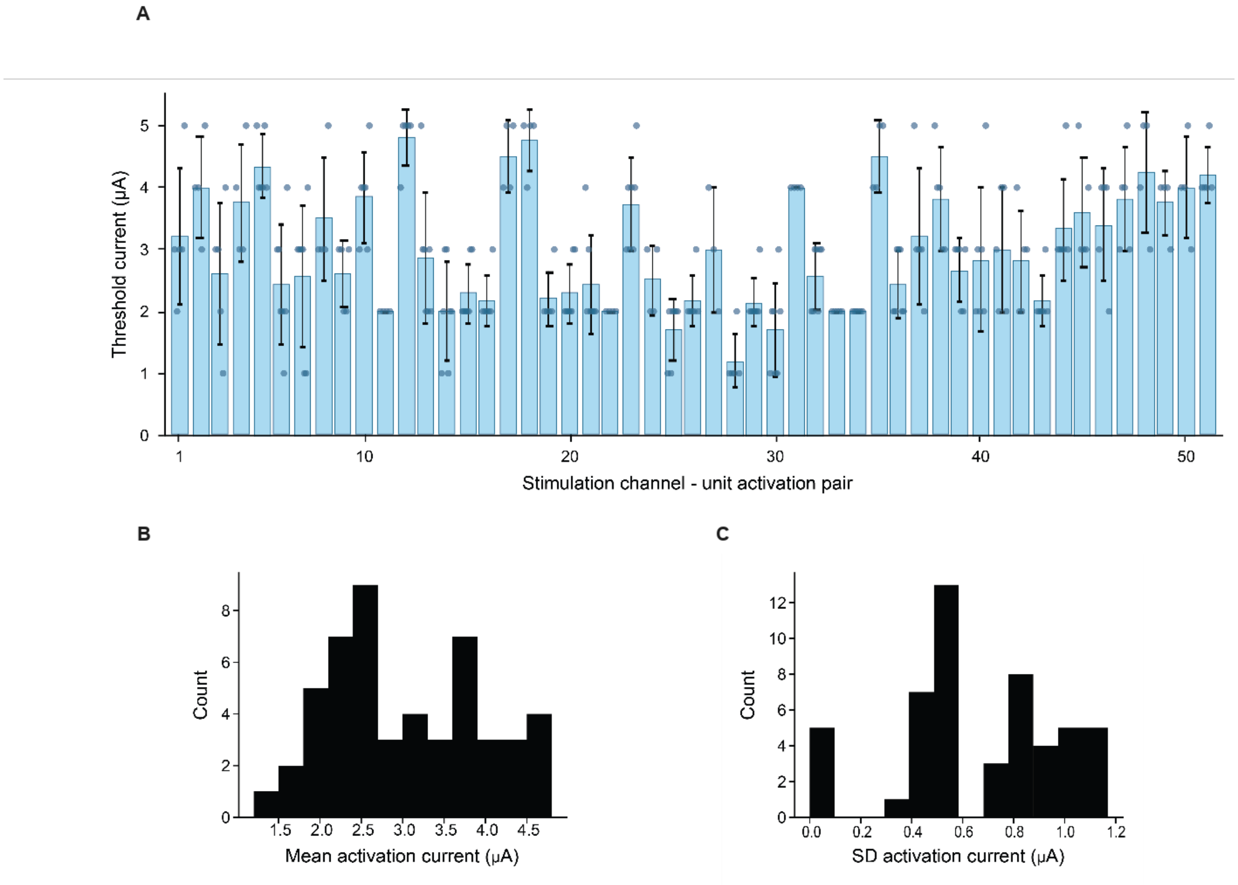


**Figure S11. Activation thresholds remain stable across longitudinal stimulation sessions.**

(A) Session-level activation threshold currents for 51 significant stimulation channel - unit pairs across seven ICMS sessions. Bars show the mean threshold for each pair and whiskers show the session-to-session SD; individual session values are shown as points. (B) Distribution of mean activation thresholds across the 51 pairs. Mean thresholds ranged from 1.2 to 4.8 μA, with a median of 2.8 μA and a mean of 3.0 μA. (C) Session-to-session threshold variability for each unit-stimulation channel pair. Threshold SDs were modest across pairs, with median SD 0.7 μA, mean SD 0.8 μA, and maximum SD 1.2 μA, indicating that activation thresholds were largely stable over repeated sessions.

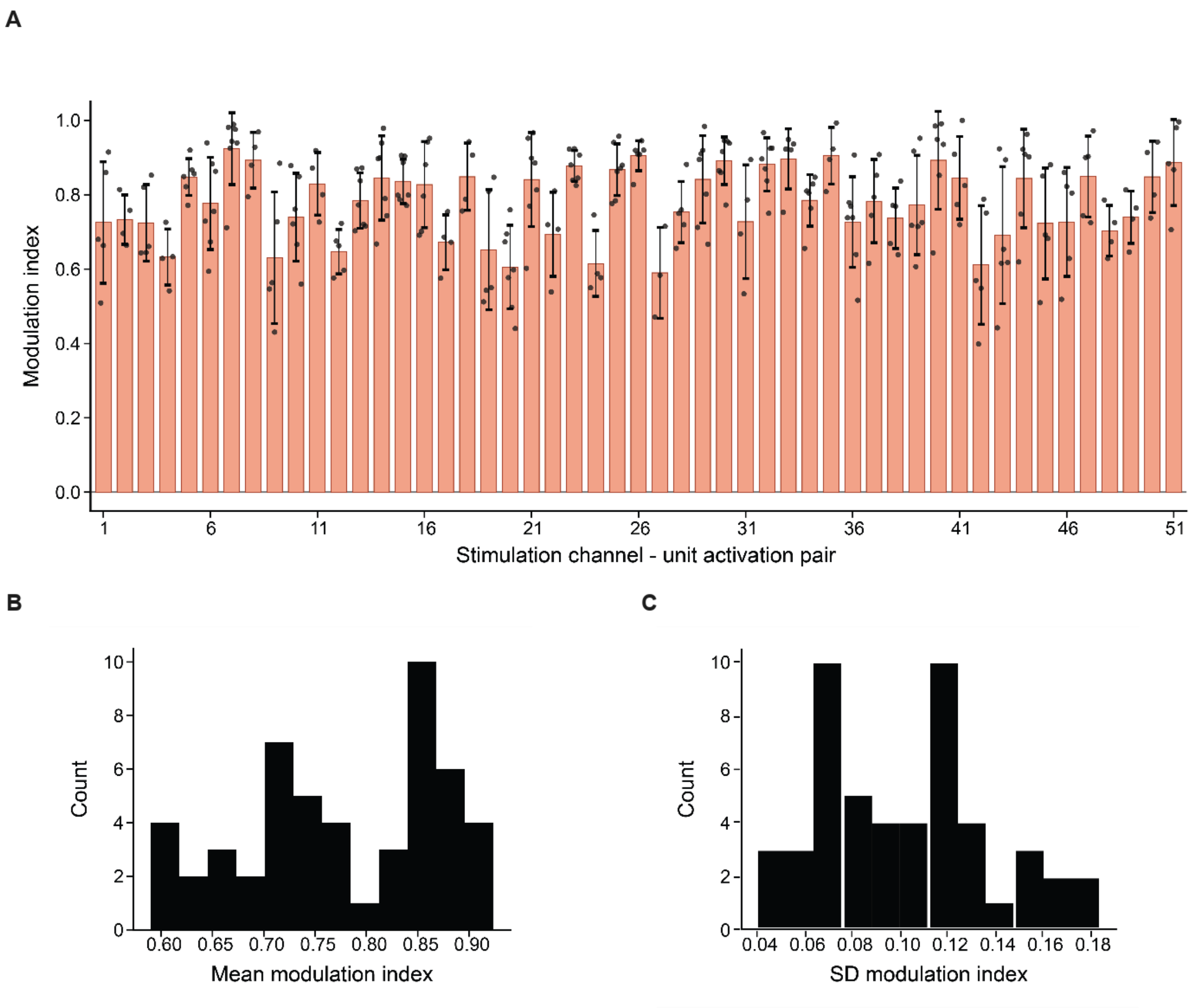


**Figure S12. Stimulation-evoked modulation is stable across sessions.** (A) Session-level stimulation modulation for the same 51 channel – unit pairs. Bars show the mean modulation index (STAR Methods) for each pair, whiskers show SD across sessions, and points show individual session values. (B) Distribution of mean modulation indices, ranging from 0.590 to 0.924, with median 0.783 and mean 0.777, indicating robust increases in firing during stimulation. (C) Session-to-session variability of the modulation index. SDs were below 0.2 for all pairs, ranging from 0.040 to 0.184 (median 0.103, mean 0.103), supporting stable stimulation-evoked modulation across repeated sessions.

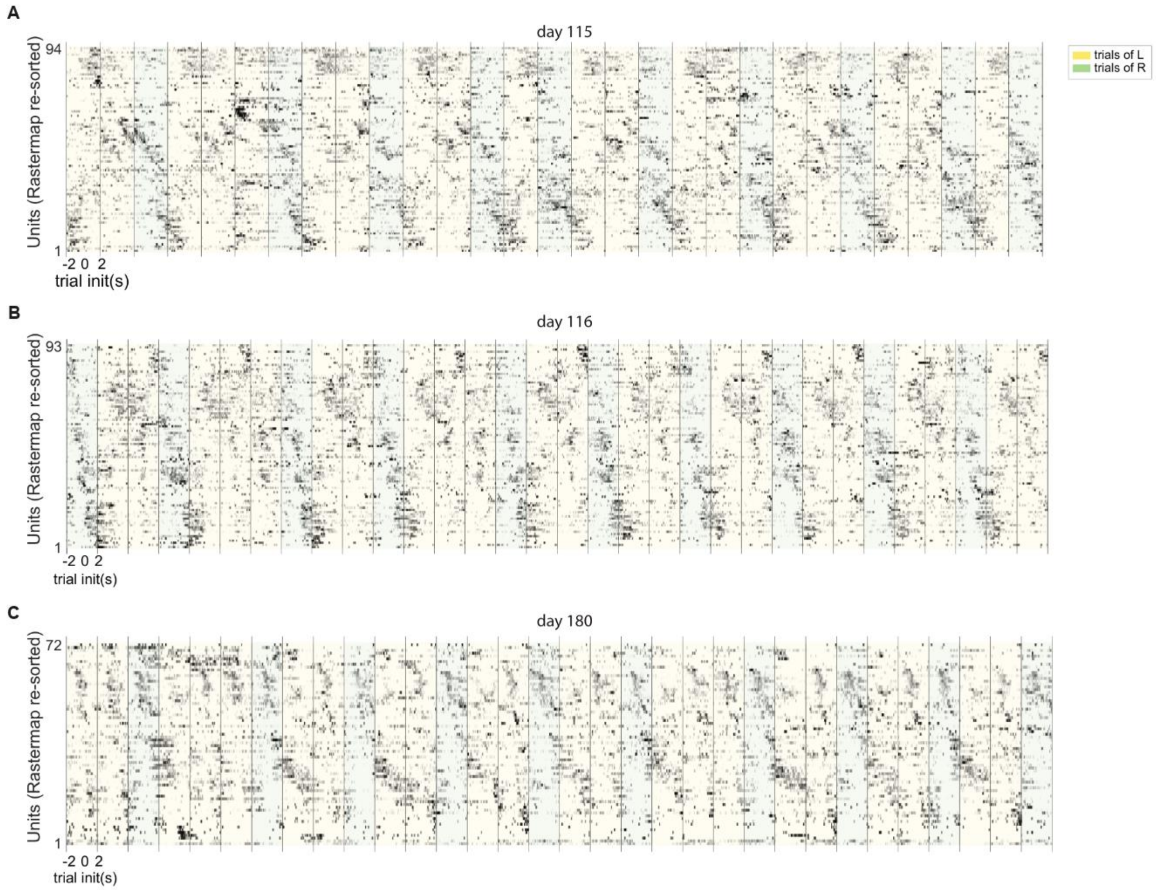


**Figure S13. Long-term recording in rat ACC during choice behavior**. Raster plots show population spiking activity during 30 consecutive trials of the learned left-left-right (L-L-R) choice sequence. Activity is shown from 2 s before to 2 s after trial initiation, with units ordered by Rastermap (as in Figure 4B). Representative recordings are shown on post-implantation days (A) 115, (B) 116, and (C) 180.

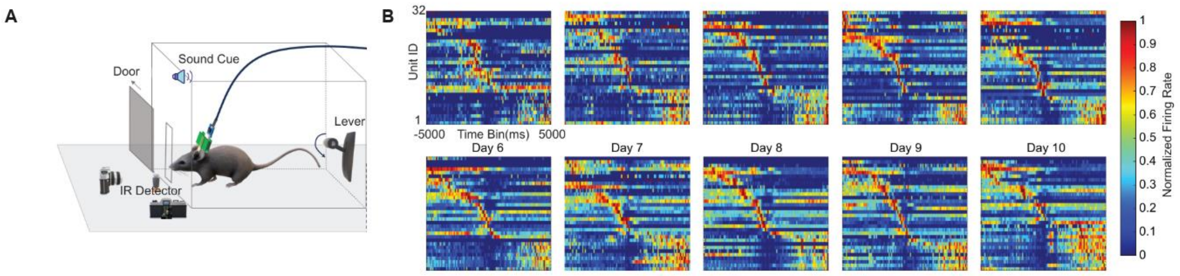


**Figure S14. Longitudinal motor-cortex activity during reach-and-grasp learning**. (A) Schematic of the single-pellet reach-and-grasp task performed by freely moving mice. Animals were trained to reach for a single-pellet reward following a sound cue. (B) Normalized firing-rate heat maps of 32 longitudinally tracked motor-cortex neurons across 10 consecutive training days, with trials aligned to the time at which the reaching paw attained its maximum distance.

Table S1: Summary of animals used in the experiments

| Animal # | Species | Location | Recording duration | Figures |
|---|---|---|---|---|
| 1 | Mouse | primary somatosensory cortex (-1.5AP, -2ML) | 6 months | Figure 2(b)(c)(d)(e)(f)(g)(h) / Figure 3(a)(b)(c)(d)(e)(f)(g) / Supplementary figure 5, 6, 8, 9, 10, 11, 12 |
| 2 | Mouse | primary somatosensory cortex (-1.5AP, -2ML) | 6 months | Figure 2(b)(c) / Figure 3(e) / Supplementary figure 8, 9 |
| 3 | Mouse | primary somatosensory cortex (-1.5AP, -1.5ML) | 6 months | Figure 2(b)(c) / Figure 3(e) / Supplementary figure 8, 9 |
| 4 | Mouse | primary somatosensory cortex (-1.5AP, -2ML) | 6 months | Figure 2(b)(c) / Figure 3(e) / Supplementary figure 8, 9 |
| 5 | Mouse | primary somatosensory cortex (-1.5AP, -2ML) | 4 months | Figure 2(b)(c) / Figure 3(e) / Supplementary figure 8, 9 |
| 6 | Mouse | primary motor cortex and | 2 months | Supplementary figure 14 |

| | | | | |
|---|---|---|---|---|
| | | dorsal striatum (0AP, 1.1ML) | | |
| 7 | Rat | Secondary motor cortex and cingulate cortex area 1 (3.0AP, -0.6ML, 2.5DV) | 6 months | Figure 2(a) / Figure 4(a)(b)(c)(d)(e) / Supplementary figure 13 |
| 8 | Rat | Hippocampus and dentate gyrus (-4.0 AP, -2.0ML, 4.5DV) | 5 months | Figure 4(f)(g)(h) |
